# Lanifibranor (IVA-337) - a pan-PPAR agonist suppresses TGF-β_1_-induced cardiac fibrosis and rescues cardiomyocyte function

**DOI:** 10.64898/2026.08.18.745414

**Authors:** Milena Paw, Lukas Minder, Andrea Laimbacher, Patrycja Kaczara, Magdalena Czepiec, Sylwia Bobis-Wozowicz, Dawid Wnuk, Barbara Kutryb-Zając, Alicja Braczko, Michał Sarna, Stefan Chłopicki, Zbigniew Madeja, Oliver Distler, Przemysław Błyszczuk, Jarosław Czyż, Gabriela Kania

**Affiliations:** Center of Experimental Rheumatology, Department of Rheumatology, University Hospital Zurich, University of Zurich, Wagistr. 14, 8952 Schlieren, Switzerland; Department of Cell Biology, Faculty of Biochemistry, Biophysics and Biotechnology, Jagiellonian University, 30-387 Cracow, Poland; Jagiellonian Centre for Experimental Therapeutics, Jagiellonian University, Bobrzynskiego 14, 30-348, Cracow, Poland; Department of Clinical Immunology, Jagiellonian University Medical College, 30-663 Cracow, Poland; Department of Biochemistry, Medical University of Gdansk, 80-211 Gdansk, Poland; Department of Biophysics, Faculty of Biochemistry, Biophysics and Biotechnology, Jagiellonian University, 30-387 Cracow, Poland

**Keywords:** cardiac microtissues, TGF-β_1_, lanifibranor, myofibroblasts, fibrosis, cardiomyocytes

## Abstract

**Background:** Cardiac fibrosis is a hallmark of many cardiovascular diseases, driven by sustained fibroblast activation and excessive extracellular matrix deposition, leading to myocardial stiffening and impaired contractility. Current therapies inadequately address this process. This study evaluated the antifibrotic potential of lanifibranor, a balanced pan-peroxisome proliferator-activated receptors (PPARs) agonist, in TGF-β_1_-induced cardiac fibrosis.

**Methods:** Human cardiac microtissues, along with 2D and 3D cardiac fibroblast and cardiomyocyte cultures, were used to assess cell viability, structure, metabolism, contractility, and gene expression.

**Results:** Lanifibranor reduced TGF-β_1_-induced fibrosis by limiting fibroblast activation and matrix deposition without affecting viability. In fibroblasts, these effects were associated with partial restoration of mitochondrial respiration and reduced focal adhesion maturation. In cardiac microtissues, lanifibranor improved contraction kinetics, decreased profibrotic transcriptional activity, and preserved bioenergetic homeostasis despite altered nucleotide balance. In cardiomyocytes, treatment normalized contractility and calcium handling while maintaining metabolic stability.

**Conclusions:** Lanifibranor attenuates TGF-β_1_-driven cardiac fibrosis by combining antifibrotic effects with metabolic and functional improvements in human models.

**Research highlights:**

- Lanifibranor reduces TGF-β_1_-induced cardiac fibrosis in human 2D/3D in vitro models.
- Fibroblast activation and ECM deposition induced by TGF-β_1_ decline with lanifibranor.
- Lanifibranor partially restores mitochondrial function in TGF-β_1_-treated fibroblasts.
- Contractility and cellular bioenergetics recover in TGF-β_1_-treated cardiomyocytes.
- Contraction recovery and metabolic rewiring occur in TGF-β_1_/lanifibranor spheroids.

**Graphical abstract:** 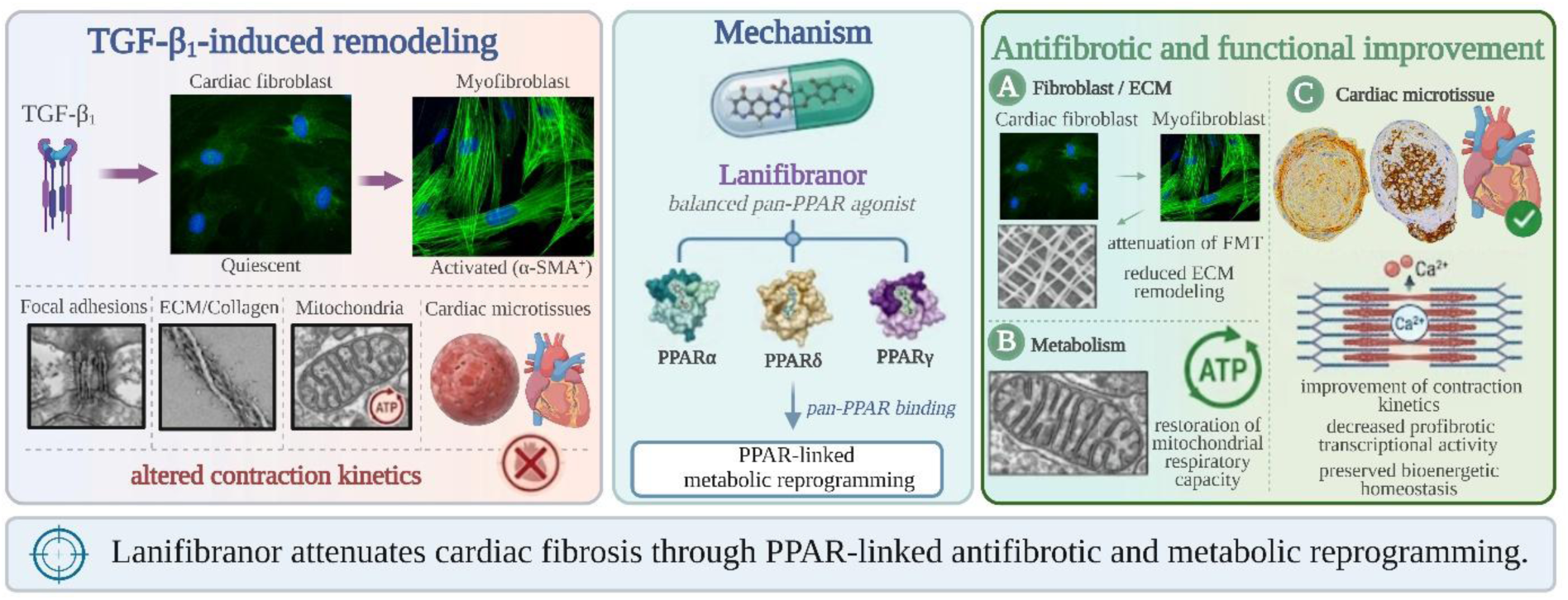

## 1. Introduction

Cardiovascular diseases (CVDs) continue to represent the foremost cause of death and disability globally, with the latest 2026 Heart and Stroke Statistics Update estimating that approximately 19.41 million global deaths in 2021 were attributable to CVD and that roughly 612 million people were living with cardiovascular disease worldwide [1]. This growing burden is driven by population ageing and the persistent prevalence of cardiometabolic risk factors, with global CVD mortality rising markedly since 1990 [2]. Across diverse etiologies of heart disease, pathological myocardial remodeling is a central determinant of progression to heart failure, and a key structural component of this remodeling is cardiac fibrosis [3]. This process is characterized by excessive deposition and remodeling of extracellular matrix (ECM) components, resulting in increased tissue stiffness and altered myocardial architecture. Importantly, fibrotic remodeling is not a passive consequence of injury but an actively regulated process that profoundly affects both the mechanical and electrical properties of the heart, impairing diastolic compliance and disrupting electrical propagation, thereby increasing vulnerability to arrhythmias [4]. Despite intensive research efforts and its well-established role as an independent predictor of morbidity and mortality across multiple forms of heart disease, there are still no approved therapies that directly and specifically reverse or prevent the progression of cardiac fibrotic remodeling in a targeted manner, underscoring a critical unmet need in cardiovascular medicine.

Cardiac fibroblasts are not a uniform population: recent single-cell and spatial transcriptomic atlases have revealed multiple fibroblast states and activation trajectories in the healthy and diseased heart, including inflammatory and ECM-remodeling subsets that emerge along a continuum rather than as a single “on/off” phenotype [5, 6]. Fibroblast activation is driven by both humoral cues and mechanical signals-soluble mediators released during injury and inflammation act together with increased matrix stiffness and cytoskeletal tension to amplify profibrotic programs [5, 7, 8]. Although TGF-β_1_ is widely recognized as the most potent inducer of fibroblast-to-myofibroblast transition, cardiac fibrosis can also be promoted by other stimuli present in cardiac disease, including angiotensin II and endothelin-1, as well as inflammatory pathways such as IL-1β-linked remodeling, highlighting that multiple profibrotic inputs can converge on a shared fibroblast effector response [9–12]. Activated fibroblasts accumulate as myofibroblasts, cells that (i) acquire contractile activity via α-smooth muscle actin-positive stress fibers and (ii) overproduce extracellular matrix proteins that are secreted and deposited in the interstitial space, progressively reshaping tissue architecture [13]. The clinical problem is not only the initiation of this response but its persistence: prolonged myofibroblast presence promotes progressively irreversible structural remodeling, increased stiffness, and ultimately functional impairment at the tissue level. At the same time, reversing established fibrosis - not merely preventing new matrix deposition - remains a major therapeutic challenge for 21st-century cardiovascular medicine, especially given the heterogeneity of fibroblast states and the multiple parallel inputs that sustain activation [5, 6].

Fibrotic remodeling of the myocardium is a central feature of heart failure progression and remains difficult to modulate pharmacologically. Growing interest has focused on the anti-fibrotic potential of peroxisome proliferator-activated receptors (PPARs), which regulate not only metabolic pathways but also inflammatory responses, mesenchymal cell differentiation, and extracellular matrix remodeling [14–17]. Clinical experience with selective PPAR agonists targeting individual isoforms, however, has revealed significant safety limitations that restricted their therapeutic use [18]. These limitations led to the development of strategies based on balanced, simultaneous activation of multiple PPAR isoforms to enhance efficacy while reducing receptor-specific exposure and improving tolerability, resulting in the emergence of a new generation of dual-and pan-PPAR agonists [19, 20]. Lanifibranor (IVA337) is a balanced pan-PPAR agonist that has shown anti-inflammatory and anti-fibrotic activity in preclinical studies, with clinical efficacy demonstrated in randomized trials in patients with metabolic liver disease [21–23]. However, its effects on cardiac fibrotic remodeling have not been systematically examined. Accordingly, this study aimed to evaluate the anti-fibrotic potential of lanifibranor using in vitro models of cardiac fibrosis, including complementary 2D and 3D culture systems and engineered cardiac microtissues [24–26].

## 2. Results

### 2.1 Lanifibranor affected the TGF-β_1_-induced myofibroblast differentiation of hCFs in 2D cultures

Using standard 2D cultures of human cardiac fibroblasts (hCFs), we evaluated cell morphology, viability, and caspase-3/7 enzymatic activity in cells cultured in reduced-serum medium (2%) and exposed to increasing concentrations of lanifibranor (Lani; 25-125 µM), in the absence or presence of TGF-β_1_ (10 ng/ml). No significant changes in cell morphology were observed at lower concentrations of lanifibranor; however, mild morphological alterations were detected at higher concentrations (100-125 µM) (Figure 1A). Although the percentage of viable fluorescein-positive cells remained unaffected up to 100 µM (Figure 1B), a dose-dependent decrease in cell metabolic activity was detected using both resazurin-based and MTT assays in hCFs treated with lanifibranor alone or in combination with TGF-β_1_ (Figure 1C-D). This reduction reflects decreased metabolic activity rather than reduced cell viability (defined as loss of cell membrane integrity). To further assess the impact of lanifibranor on cell viability, we employed a caspase-3/7 activity assay, which confirmed the lack of apoptosis induction at concentrations up to 100 µM (Figure 1E). Notably, lanifibranor significantly reduced TGF-β_1_-induced caspase-3/7 activation, restoring activity levels to those observed in control cells. In contrast, treatment with higher concentrations of lanifibranor (125 µM) resulted in increased cytotoxicity, consistent with the observed morphological alterations. Taken together, these findings indicate that lanifibranor is well tolerated by hCFs at concentrations up to 100 µM, while higher doses exert cytotoxic effects that are exacerbated in the presence of TGF-β_1_.

**Figure 1.**
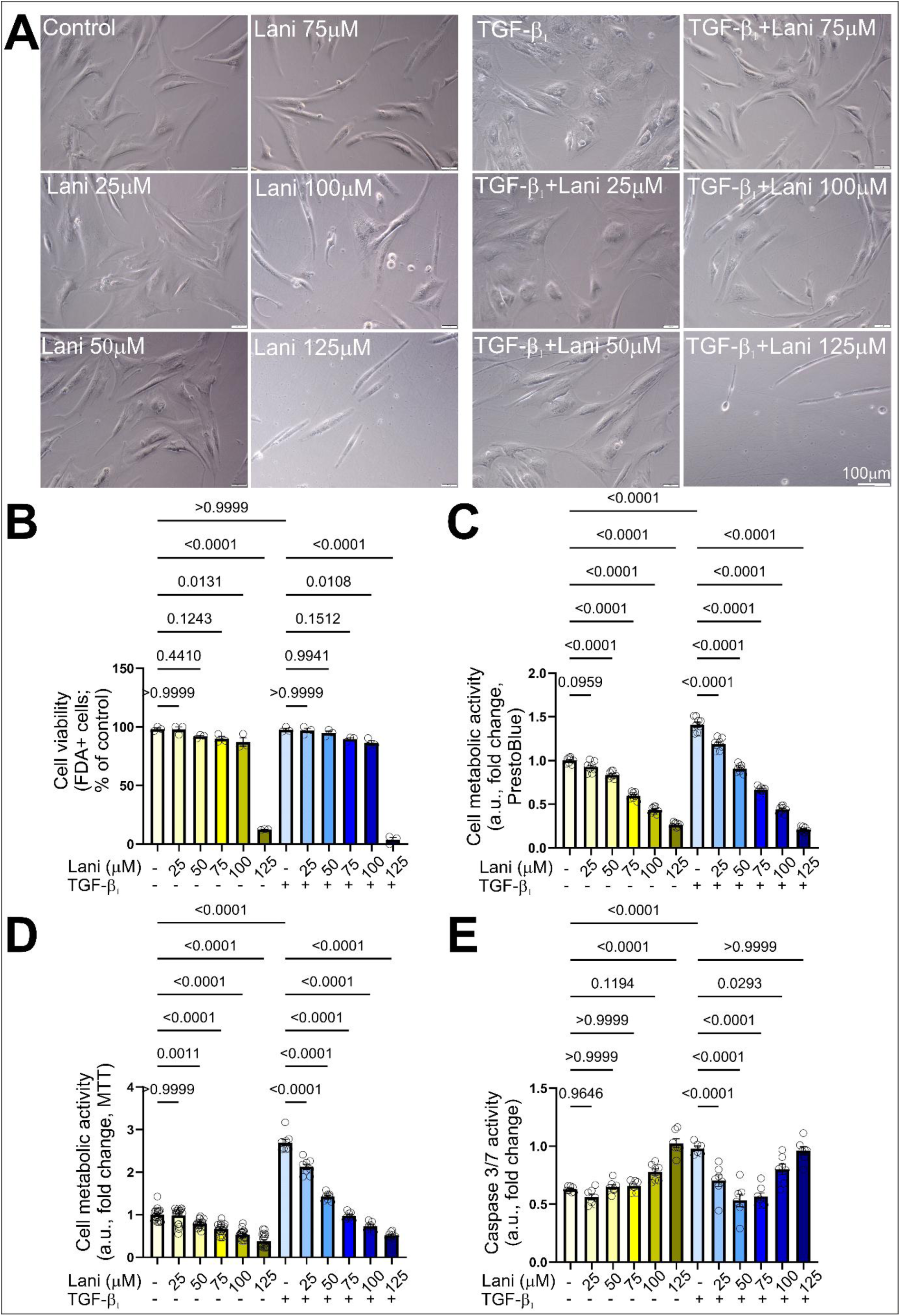
Lanifibranor did not affect cell viability or increase apoptosis, but decreased metabolic activity of hCFs. hCF cultures were treated with increasing concentrations of lanifibranor (Lani; 0 - 125 μM) in the absence or presence of TGF-β_1_ (10 ng/ml) and cultured for 4 days. **(A)** Representative images of hCFs cultured under the described conditions. Scale bar: 100 μm. Cell viability was assessed on day 4 using **(B)** FDA/EtBr staining. Cell metabolic activity was determined using **(C)** PrestoBlue and **(D)** the MTT assay. **(E)** Caspase 3/7 enzymatic activity was also measured in cultured hCFs. Data are presented as mean ± SD. Statistical significance was assessed using one-way ANOVA followed by Tukey’s post hoc test. p values are indicated in each graph.

As previously described, hCFs cultured under standard 2D *in vitro* conditions provide a robust model for analyzing the mechanisms underlying cardiac fibrosis [27]. To assess the potential involvement of lanifibranor in TGF-β_1_-induced fibroblast-to-myofibroblast transition (FMT) in hCF cultures, we investigated its effects on α-SMA and procollagen 1α1 expression levels. Lanifibranor reduced TGF-β_1_-induced α-SMA expression in a dose-dependent manner, with strong inhibition observed at concentrations ≥50 μM, and had minimal effects when applied alone at lower doses (Figure 2A). Similarly, procollagen 1α1 levels in hCFs were significantly reduced by lanifibranor in a dose-dependent manner, both in the presence and absence of TGF-β_1_ (Figure 2B). These results led us to select 50 μM lanifibranor for subsequent experiments, as this concentration did not alter morphology, viability, or caspase-3/7 activity, either alone or in combination with TGF-β_1_, and significantly attenuated the reduction in the major myofibroblast marker, α-SMA. Upon TGF-β_1_ stimulation, hCFs developed an expanded network of α-SMA-enriched microfilament bundles (Figure 2C) and promoted myofibroblastic differentiation (Figure 2E). This effect was accompanied by an increase in the size (area and length) of focal adhesions (FAs) (Figure 2D, F-H). Analysis of FA size distribution revealed that lanifibranor reduced the abundance of supermature FAs (>6 μm) in TGF-β_1_-treated hCFs (Figure 2H). When applied at a non-cytotoxic concentration (50 μM; see Figure 1A-E), lanifibranor decreased the fraction of myofibroblasts (Figure 2C, E) and attenuated FA maturation in TGF-β_1_-treated cells (Figure 2D, F-H). Given that myofibroblast differentiation is tightly associated with metabolic reprogramming [28], we next examined whether lanifibranor affects mitochondrial respiration in hCFs by measuring oxygen consumption rates (OCR) using the Seahorse XF Mito Stress Test (Figure 2I). OCR values were normalized to cell number and analyzed under basal conditions, maximal respiration, spare respiratory capacity, and ATP-linked respiration. TGF-β_1_ stimulation did not significantly alter basal respiration; however, it resulted in a marked reduction in maximal respiration and spare respiratory capacity, accompanied by a slight increase in ATP-linked respiration. Treatment of TGF-β_1_-stimulated hCFs with lanifibranor did not affect basal respiration, but partially improved maximal respiration toward control levels and markedly increased spare respiratory capacity and ATP-linked respiration. Taken together, TGF-β_1_ promotes fibroblast-to-myofibroblast differentiation accompanied by metabolic reprogramming in hCFs. Lanifibranor effectively counteracts these profibrotic effects by partially reversing TGF-β_1_-driven metabolic alterations, thereby limiting fibroblast activation and fibrotic responses in 2D hCF cultures.

**Figure 2.**
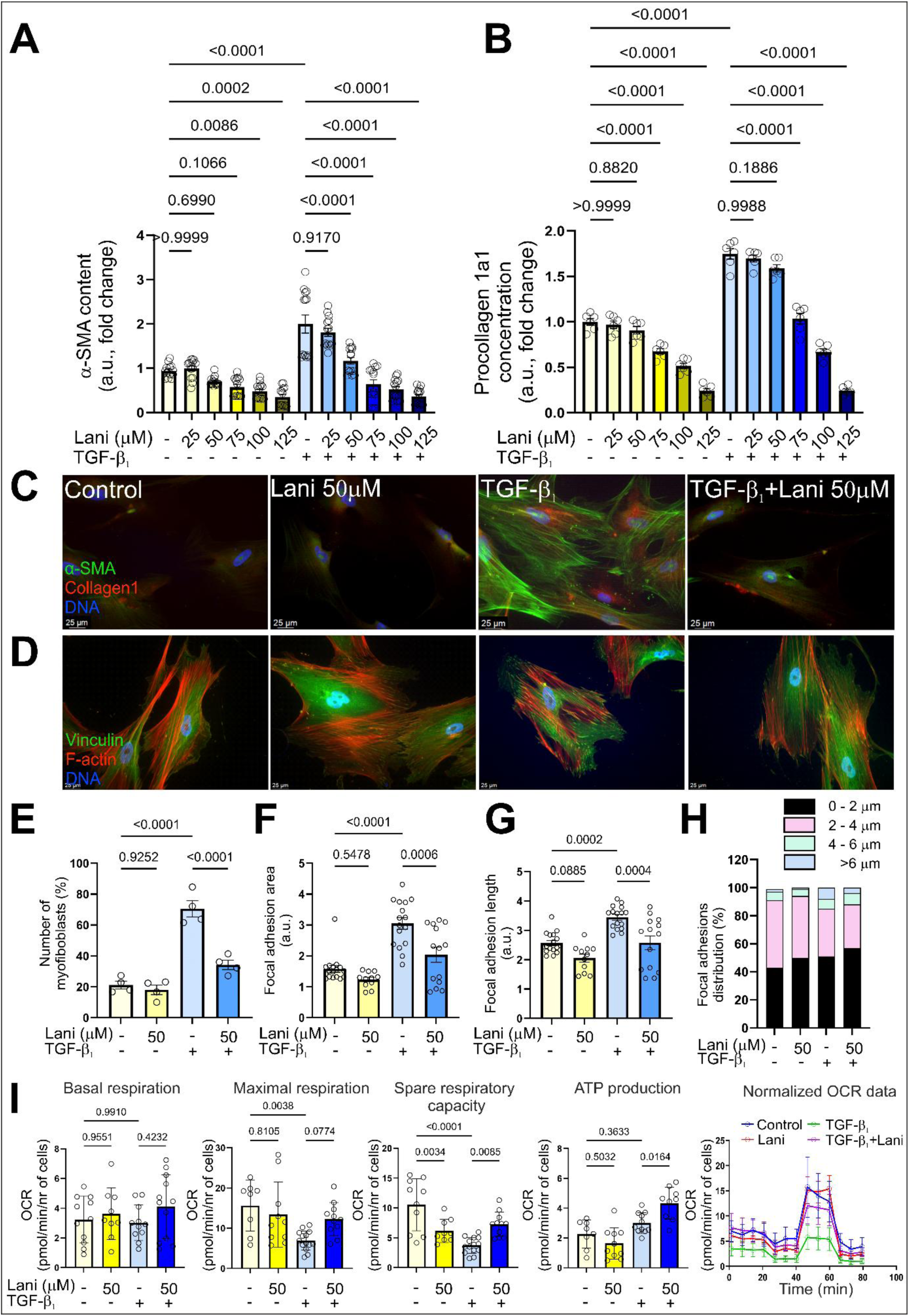
Lanifibranor suppressed TGF-β₁-induced myofibroblast transition and modulated cellular bioenergetics in hCFs. hCF were treated with lanifibranor (Lani; 0-125 μM) in the absence or presence of TGF-β_1_ (10 ng/ml) and cultured for 4 days. **(A)** α-SMA in cells and **(B)** procollagen 1α1 in supernatants were measured by ELISA. **(C)** Intracellular localization of α-SMA-positive microfilament bundles and collagen 1α1 (upper row), as well as **(D)** focal adhesion sites and actin cytoskeleton architecture (lower row), were visualized using immunofluorescence staining. Representative images are shown. Scale bars: 25 μm. **(E)** The percentage of α-SMA-positive myofibroblasts in hCF cultures was quantified**. (F)** Focal adhesion area and **(G)** length were measured from the collected images using Fiji (ImageJ) software. **(H)** Focal adhesions were categorized into defined length intervals, and their abundance was expressed as a percentage of the total number of measured contacts. **(I)** Trajectories and OCR values were normalized to cell number for all tested conditions. Cellular bioenergetic changes are presented as basal respiration, maximal respiration, spare respiratory capacity and ATP-linked respiration. Data are presented as mean ± SD. Statistical significance was assessed using one-way ANOVA with Tukey’s post hoc test or the Kruskal–Wallis test with Dunn’s post hoc test, as appropriate. p values are indicated in each graph.

### 2.2 Lanifibranor did not alter the viability or increase apoptosis, but significantly reduced procollagen 1a1 level in TGF-β_1_-treated 3D spheroid hCF cultures

To specifically assess lanifibranor in a fibrosis-relevant 3D microenvironment, we established a 3D spheroid model of human cardiac fibroblasts (hCFs), which enables the capture of key cell-cell/ECM interactions and remodeling dynamics that are difficult to reproduce in 2D [29]. Using this 3D hCF spheroid model, we evaluated the effects of lanifibranor on cell viability, apoptosis, and procollagen 1α1 production. In accordance with previous findings [26], 3D models required higher concentrations of TGF-β_1_ than 2D cultures due to restricted diffusion. Compact, round fibroblast spheroids were maintained following treatment with increasing concentrations of lanifibranor (Lani; 0-125 μM), both in the absence and presence of TGF-β_1_ stimulation (Figure 3A). A resazurin-based metabolic activity assay demonstrated that lanifibranor, at concentrations up to 100 μM, did not impair the relative cell viability (based on metabolic activity) of hCF spheroids cultured with or without TGF-β_1_ (Figure 3B). Furthermore, no significant changes in caspase 3/7 activation were observed in spheroids treated with lanifibranor alone or in combination with TGF-β_1_ at concentrations of 50 or 75 μM (Figure 3C). Notably, the elevated levels of procollagen 1α1 induced by TGF-β_1_ in spheroid cultures were significantly reduced by lanifibranor at concentrations above 25 μM (Figure 3D). These findings demonstrate for the first time that lanifibranor at 50 and 75 μM attenuates TGF-β_1_-induced fibroblast activation without adversely affecting cell viability in 3D spheroid cultures.

**Figure 3.**
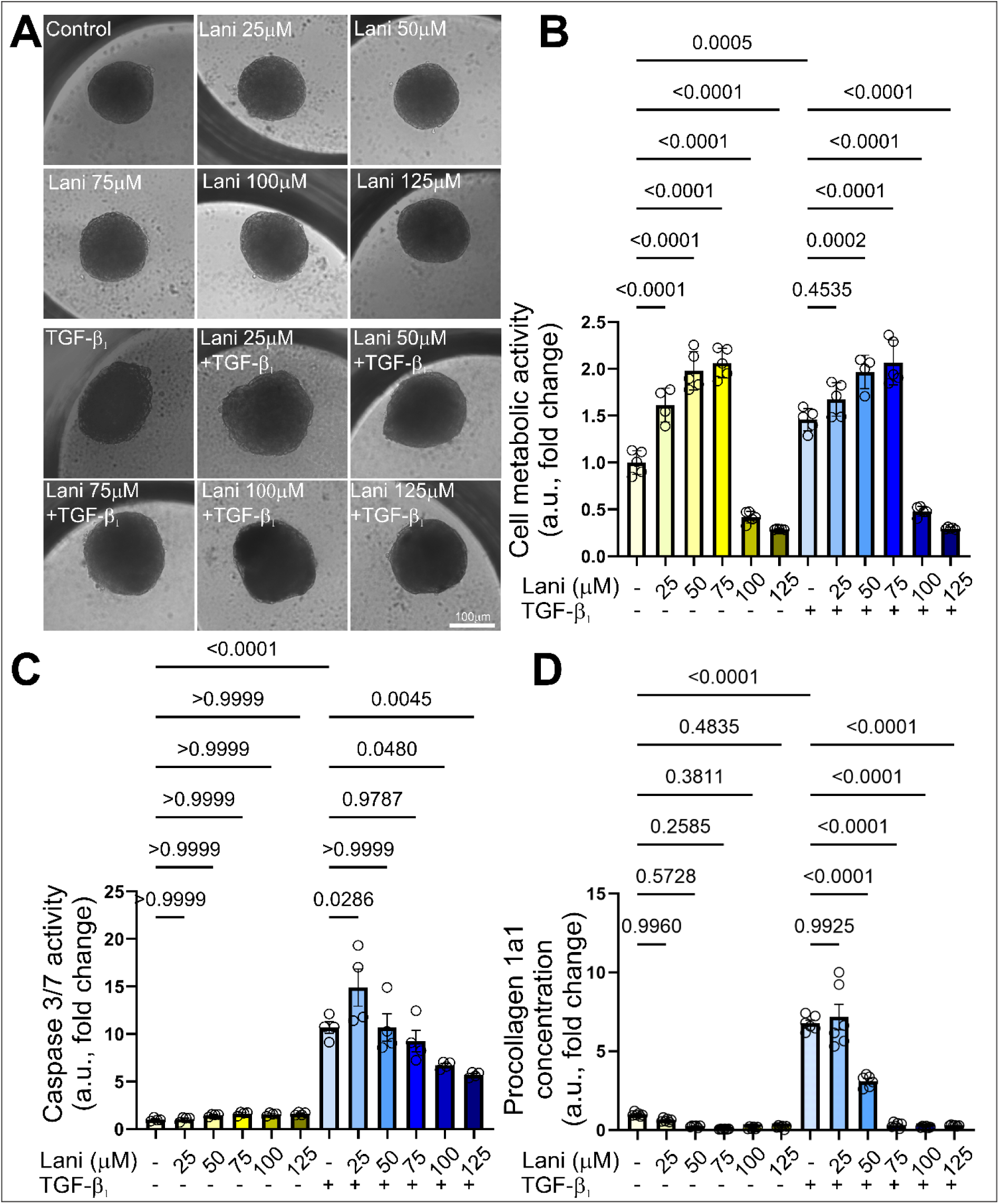
Lanifibranor did not affect cell viability and attenuates TGF-β₁-induced procollagen 1a1 secretion in three-dimensional (3D) cultures of hCFs. Cells were treated with increasing concentrations of lanifibranor (Lani; 0-125 μM) in the absence or presence of TGF-β_1_ (20 ng/ml) and cultured for 10 days. **(A)** Representative images of hCF 3D cultures. **(B)** Metabolic activity (used as an indicator of viability) was assessed using the PrestoBlue HS reagent. **(C)** Caspase 3/7 activity and **(D)** concentration of secreted procollagen 1α1 in hCF 3D cultures on day 10 were measured. Data are presented as mean ± SD. Statistical significance was assessed using one-way ANOVA followed by Tukey’s post hoc test. p values are indicated in each graph.

### 2.3 Lanifibranor did not alter cell viability or apoptosis, but significantly reduced fibrosis in cardiac microtissues

As previously reported, to better model cardiac function during the development of cardiomyopathy, we used an *in vitro* model of human cardiac microtissues composed of human iPSC-derived cardiomyocytes (hiPSC-CMs) and human fetal cardiac fibroblasts (hCFs) [26]. Based on our initial results, we selected 50 μM lanifibranor concentration for subsequent microtissue experiments, as it did not alter morphology or viability and had no effect on caspase-3/7 activity, either alone or with TGF-β_1_ (Figure 4A-C). Overall, cell survival remained unaffected, indicating that this dose is well tolerated in 3D cardiac microtissue cultures under the experimental conditions.

**Figure 4.**
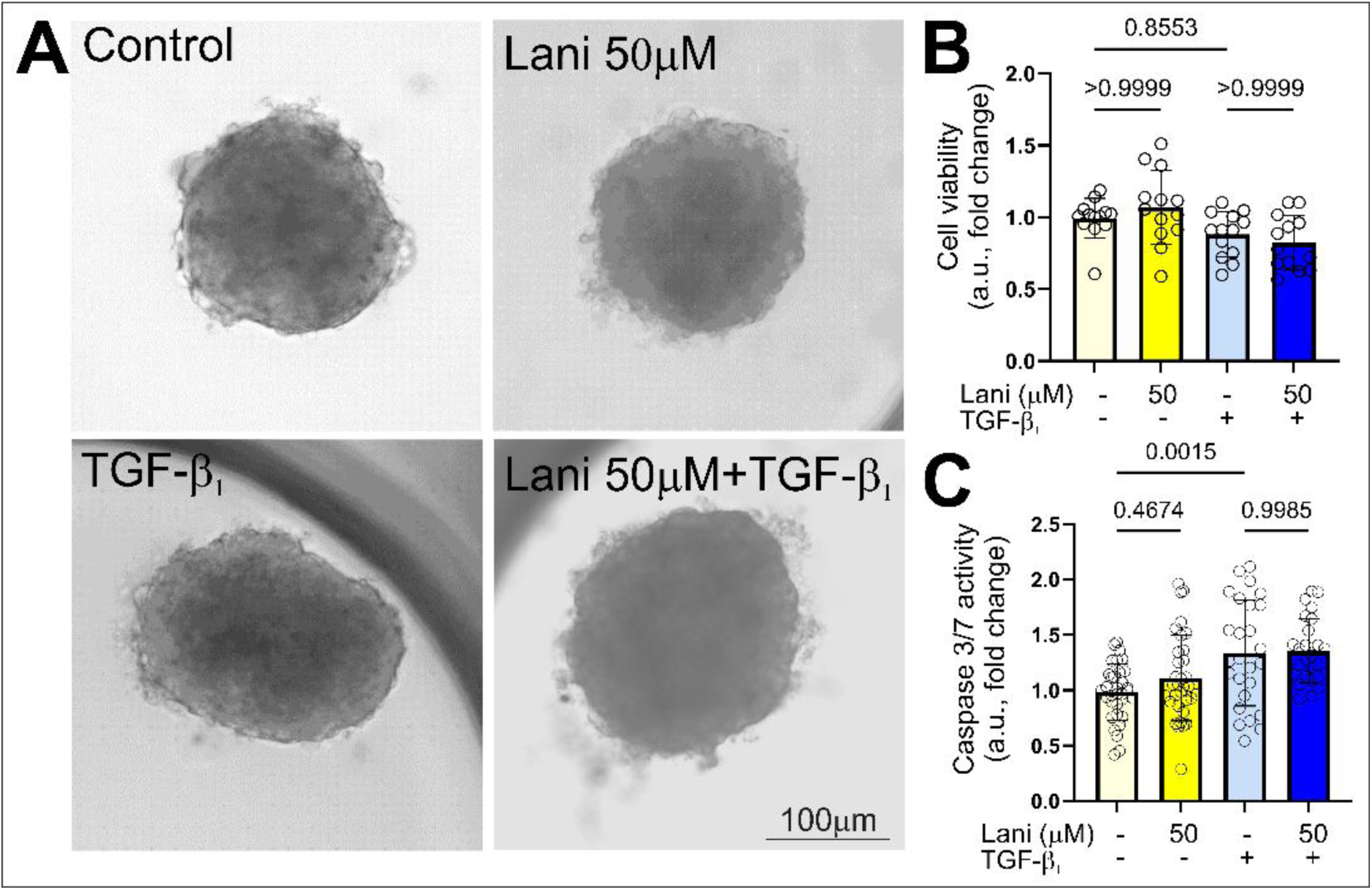
Lanifibranor did not affect the viability or apoptosis of TGF-β_1_-stimulated cardiac microtissues. Cardiac microtissues (MTs) treated with lanifibranor (Lani; 50 μM) in the absence or presence of TGF-β_1_ (20 ng/ml) and cultured for 10 days. **(A)** Representative images of MTs were presented. **(B)** Caspase 3/7 activity and **(C)** cell viability were measured in cultures of MTs at day 10. Data are presented as mean ± SD. Statistical significance was tested using the one-way ANOVA with Tukey’s post-hoc test. p values are presented in each graph.

Global transcriptomic profiling was performed to characterize and compare the molecular effects of lanifibranor and TGF-β₁ administered individually or in combination in cardiac microtissues. Differential gene expression analysis was conducted using a threshold of |log2 fold change| ≥ 1 and an adjusted p-value < 0.05. Assessment of overlap and specificity using UpSet analysis and Venn diagrams showed that most upregulated and downregulated DEGs were contrast-specific, with limited overlap between comparisons (Figure S1, 5A), a pattern further illustrated by volcano plots (Figure 5C). Hierarchical clustering of the union of differentially expressed genes (DEGs) demonstrated distinct transcriptional profiles across all experimental conditions (Figure 5B). Collectively, these analyses indicate that lanifibranor induces a distinct, context-dependent transcriptional response and selectively modulates TGF-β₁-regulated gene expression rather than eliciting uniform global changes.

**Figure 5.**
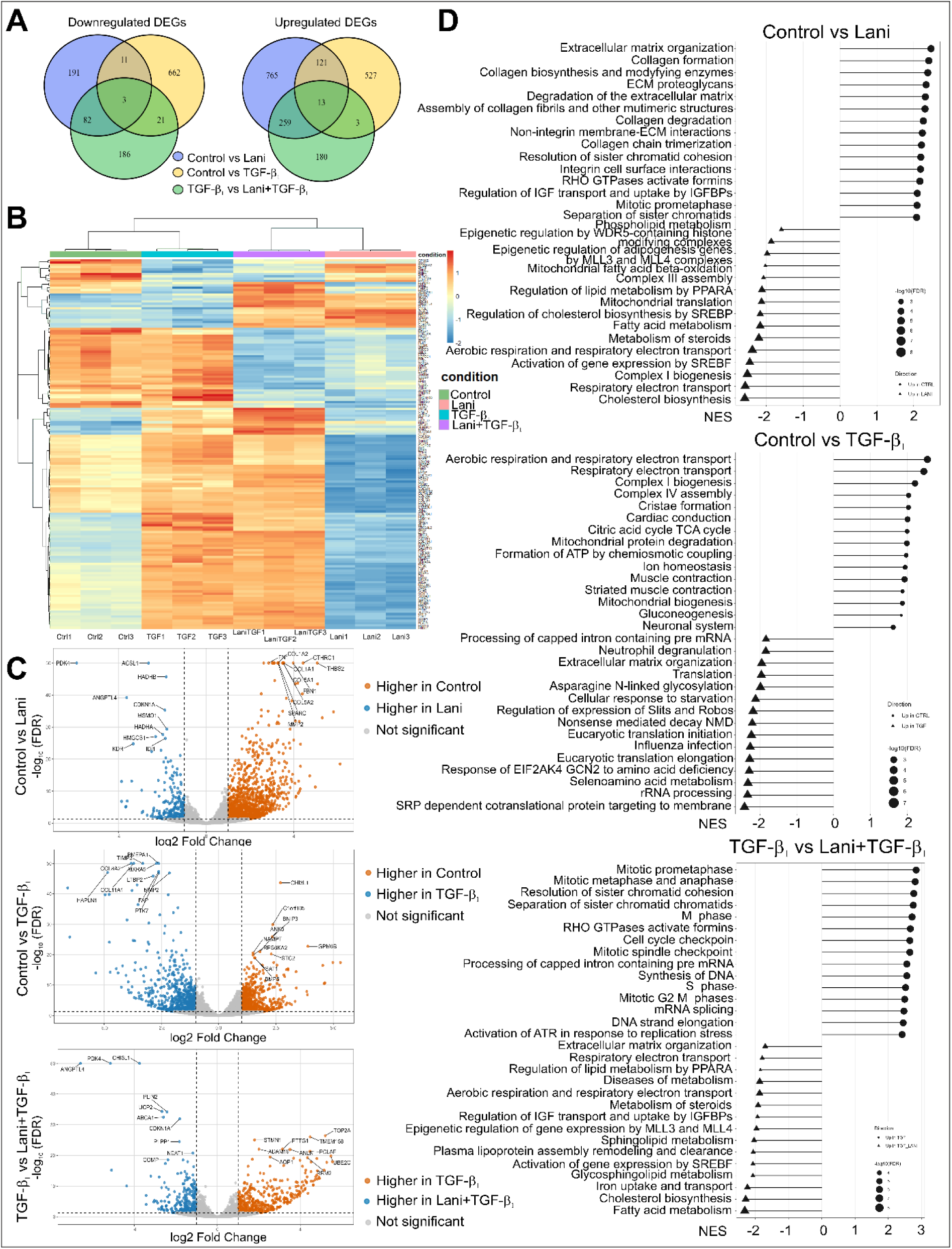
Global transcriptomic profiling reveals lanifibranor-dependent modulation of TGF-β₁-driven gene expression in cardiac microtissues. Transcriptional profiling and functional analysis of differential gene expression were performed using MTs cultured in the absence or presence of TGF-β₁ (20 ng/ml) without or with lanifibranor (50 μM) for 10 days. **(A)** Venn diagrams illustrating shared and unique DEGs among pairwise comparisons. **(B)** Heatmap of DEGs (|log2FC| ≥ 1, adjusted p-value < 0.05) across all comparisons with hierarchical clustering of genes and samples. **(C)** Volcano plots showing gene expression fold changes and statistical significance for each comparison. Differentially expressed genes were defined as |log₂FC| ≥ 1 and adjusted p-value < 0.05; the top 10 upregulated and top 10 downregulated DEGs are highlighted. **(D)** Reactome pathway enrichment analysis based on gene set enrichment analysis (GSEA), revealing contrast-dependent enrichment of pathways related to lipid and energy metabolism, mitochondrial respiration, extracellular matrix organization, cell cycle processes, and PPAR-related signaling.

Functional enrichment analysis using Reactome pathway annotation revealed distinct and condition-specific transcriptional programs (Figure 5D, S1). Compared with control, lanifibranor-treated microtissues displayed a metabolic transcriptional shift. GSEA/Reactome analysis showed enrichment of pathways related to fatty acid metabolism, mitochondrial β-oxidation, lipid metabolism, cholesterol biosynthesis, steroid metabolism, and respiratory electron transport (Figure 5D). Consistent with this, DEG analysis revealed upregulation of metabolic genes, including *PDK4*, *ACSL1*, *HADHB*, *ANGPTL4*, *HADHA*, *HMGCS1*, and *IDI1* (Figure 5C, Table S1). Compared with control, TGF-β₁-treated microtissues showed enrichment of pathways related primarily to extracellular matrix organization, as well as translation-, glycosylation-, and RNA-processing-associated processes (Figure 5D). Consistent with this, DEG analysis revealed increased expression of matrix remodeling-and TGF-β-associated genes, including *TIMP3, COL11A1, COL8A2, PMEPA1, MXRA5, FAP, HAPLN1, MMP2, PTK7,* and *LTBP2* (Figure 5C, Table S2). Compared with TGF-β₁ alone, the addition of lanifibranor partially redirected the TGF-β₁-driven transcriptional response. Reactome GSEA indicated enrichment of metabolic pathways in the TGF-β₁ + lanifibranor group, including fatty acid metabolism, cholesterol biosynthesis, steroid metabolism, respiratory electron transport, and lipid metabolism by PPARA, whereas TGF-β₁ alone remained associated mainly with cell cycle-and mitosis-related pathways (Figure 5D). In agreement with this, DEG analysis showed increased expression in the TGF-β₁ + lanifibranor group of genes associated with metabolic regulation, such as PDK4, ANGPTL4, UCP2, PLIN2, ABCA1, CDKN1A, and PLPP1 (Figure 5C, Table S3). Complementary analyses of DEG overlap across comparisons and Reactome over-representation analysis (ORA) performed on DEG sets are provided in Supplementary Figure S1. The top 10 upregulated and top 10 downregulated DEGs for each contrast are listed in Supplementary Tables S1-S3.

Overall, these data demonstrate that TGF-β₁ induces a broad profibrotic transcriptional program in cardiac microtissues, whereas lanifibranor promotes metabolic and mitochondrial transcriptional signatures and partially redirects TGF-β₁-associated transcriptional responses when administered in combination.

Based on transcriptomic evidence indicating lanifibranor-dependent modulation of metabolic and ECM gene expression programs, we next assessed whether these changes translated into structural alterations in cardiac microtissues using immunohistochemistry. TGF-β_1_ stimulation markedly increased the expression of fibroblast activation markers α-SMA, fibronectin, and periostin, consistent with myofibroblast differentiation (Figure 6A). Quantitative image analysis demonstrated that co-treatment with lanifibranor significantly reduced the levels of these fibrosis-associated markers (Figure 6B-D). In parallel, TGF-β_1_-induced upregulation of profibrotic markers was accompanied by reduced expression of the cardiomyocyte marker troponin T, and this effect was not prevented by lanifibranor (Figure 6A, E). Masson’s Trichrome staining revealed increased collagen accumulation in TGF-β_1_-treated microtissues, which was significantly attenuated by lanifibranor (Figure 6A, F). Consistently, lanifibranor treatment reduced TGF-β_1_-enhanced secretion of procollagen 1a1 in microtissues (Figure 6G). Collectively, these data indicate that lanifibranor attenuates TGF-β_1_-induced fibroblast activation and ECM remodeling in 3D cardiac microtissues.

**Figure 6.**
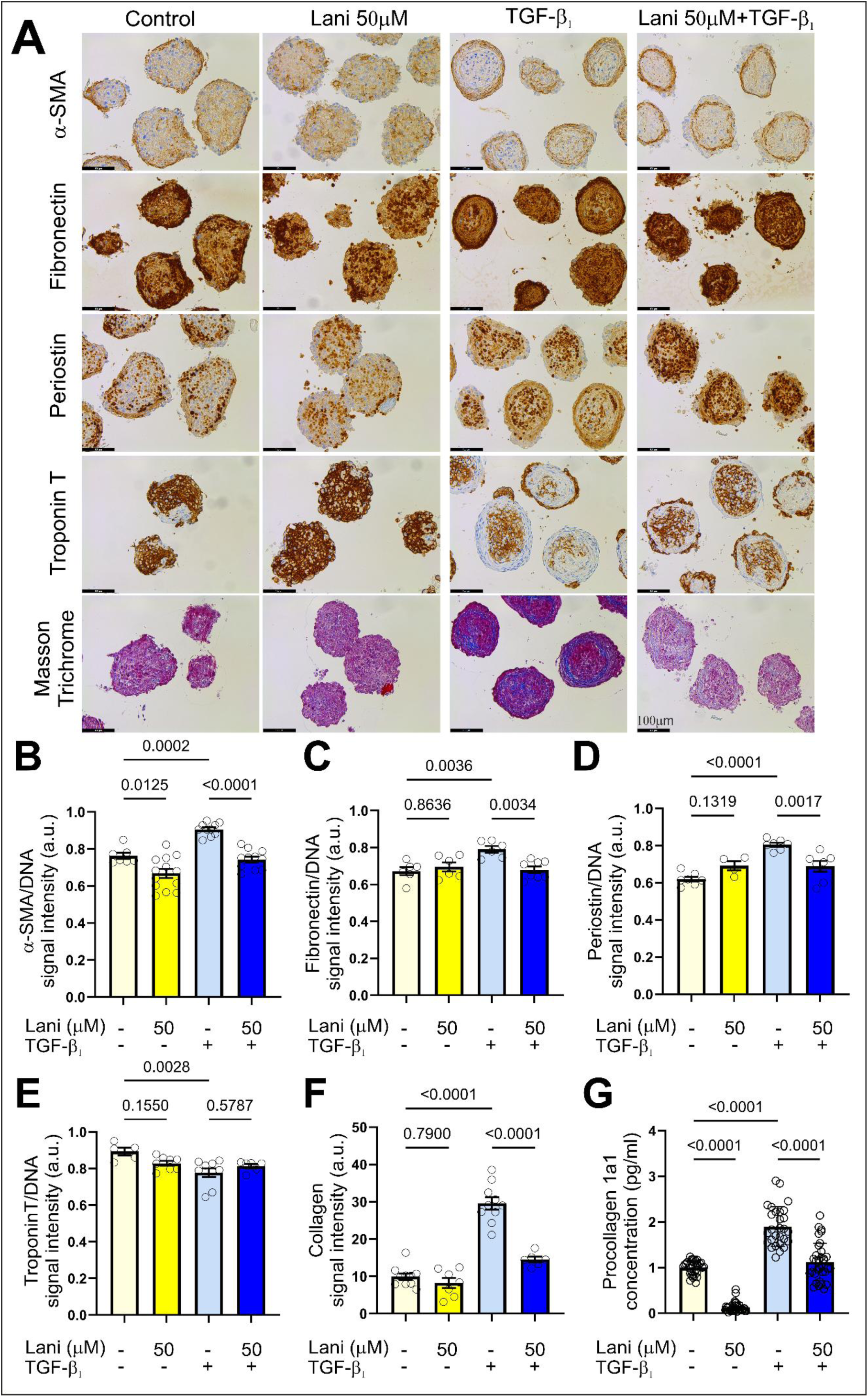
Lanifibranor limits TGF-β_1_-driven fibrosis in cardiac microtissues. Immunohistochemical analysis was performed using cardiac microtissues (MTs) treated with lanifibranor (Lani; 50 μM) in the absence or presence of TGF-β_1_ (20 ng/ml) and cultured for 10 days. **(A)** Representative images of MTs immunostained for α-SMA, fibronectin, periostin, Masson’s Trichrome, and troponin T. Scale bar = 100 μm. Quantitative analysis of immunostained MTs for **(B)** α-SMA, **(C)** fibronectin, **(D)** periostin, **(E)** troponin T, and **(F)** Masson’s Trichrome content. **(G)** Procollagen 1a1 concentration in supernatants collected from MT cultures on day 10 (n = 30). Data are presented as mean ± SD. Statistical significance was assessed using one-way ANOVA followed by Tukey’s post hoc test. p values are indicated in each graph.

### 2.4 Lanifibranor modulated the contraction properties of TGF-β_1_-treated cardiac microtissues without affecting cellular bioenergetics

We next examined whether lanifibranor-dependent transcriptional and structural changes translated into functional alterations in cardiac microtissue contractility. Using high-speed video recordings and motion-tracking analysis [30, 31], we quantified key contractile parameters in microtissues cultured with or without TGF-β_1_ and lanifibranor. TGF-β_1_ stimulation significantly reduced contraction duration (Figure 7A), contraction amplitude (Figure 7B), time-to-peak (Figure 7C), and relaxation time (Figure 7D), while increasing beating rate (Figure 7E). Lanifibranor (50 μM) effectively restored contraction duration, time-to-peak, and relaxation time to near-control levels and significantly attenuated the TGF-β_1_-induced increase in beating rate but did not rescue contraction amplitude in TGF-β_1_-stimulated microtissues.

**Figure 7.**
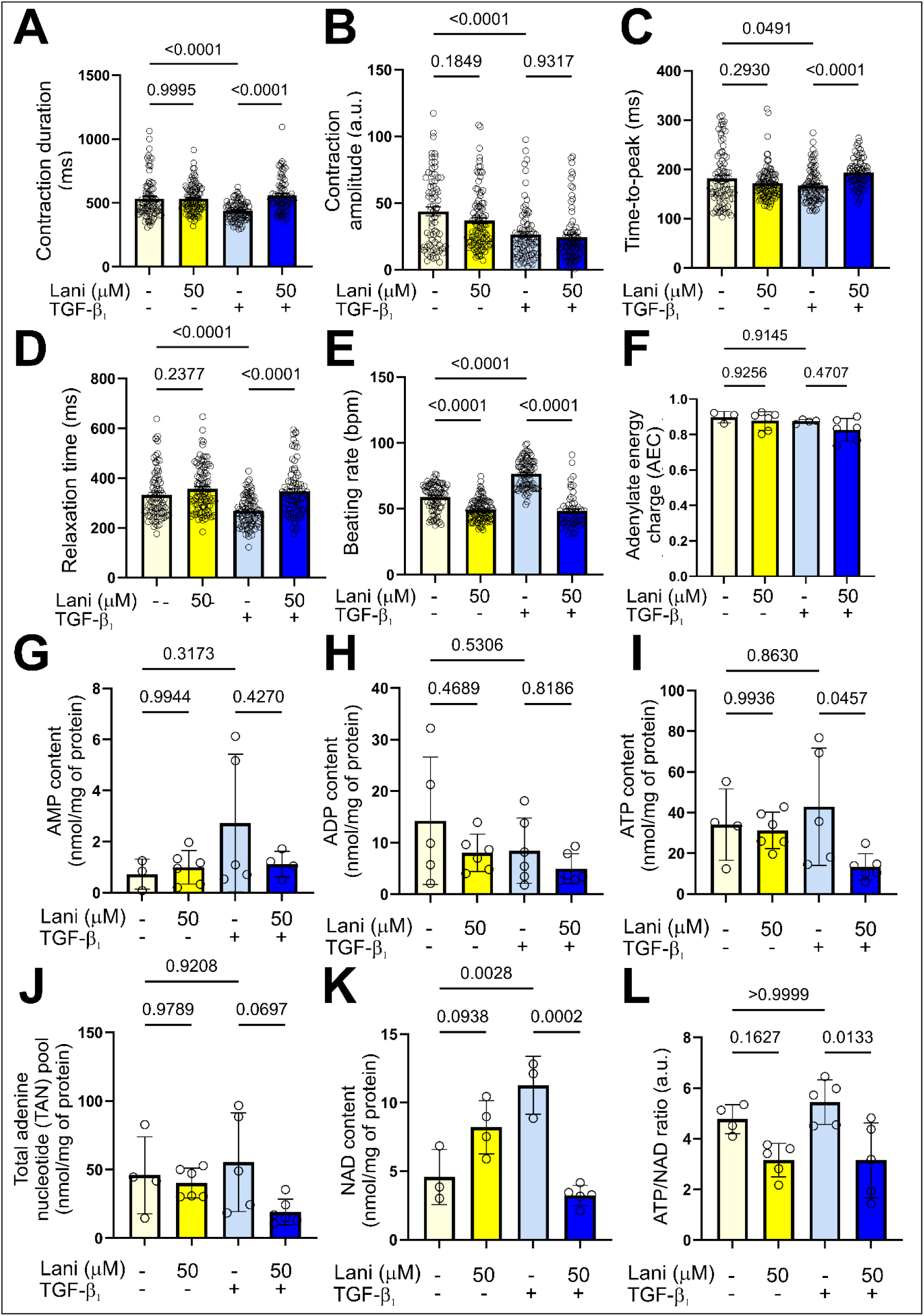
Lanifibranor modulates contractile function and nucleotide balance in TGF-β_1_-stimulated cardiac microtissues. Contractile properties and nucleotide concentrations were determined in cardiac microtissues (MTs) treated with lanifibranor (Lani; 50 μM) in the absence or presence of TGF-β_1_ (20 ng/ml) and cultured for 10 days. Contractions of MTs were recorded for 15 seconds using a Zeiss Axio Observer microscope. Quantification of contractile parameters was performed using the MUSCLEMOTION macro and Fiji (ImageJ) software. The parameters analyzed included: **(A)** contraction duration, **(B)** contraction amplitude, **(C)** time-to-peak, **(D)** relaxation time and **(E)** beating rate. Each dot represents data from one MT. At least 15 MTs were analyzed per condition in each of three independent experiments. Nucleotide concentrations were measured using ultra-high-performance liquid chromatography with diode array detection (UHPLC-DAD). **(F)** Adenylate energy charge and the content of **(G)** AMP, **(H)** ADP, **(I)** ATP, **(J)** the total adenine nucleotide (TAN) pool, **(K)** NAD content, and **(L)** the ATP/NAD ratio was quantified. Values were normalized to total protein content and are presented as mean ± SD (n ≥ 5). Statistical significance was assessed using one-way ANOVA followed by Tukey’s post hoc test; p values are indicated in the graphs.

To assess whether these functional effects were associated with altered cellular energy status, nucleotide metabolism was analyzed by UHPLC-DAD. Adenylate energy charge (AEC) did not differ significantly between conditions, consistent with maintained bioenergetic homeostasis (Figure 7F). TGF-β₁ stimulation alone did not significantly affect AMP, ADP, ATP, or total adenine nucleotide (TAN) levels (Figure 7G-J) or the ATP/NAD ratio (Figure 7L). However, TGF-β₁ significantly increased NAD content (Figure 7K). In contrast, lanifibranor treatment of TGF-β_1_-stimulated microtissues significantly reduced ATP levels (Figure 7I), with only modest changes in AMP, ADP, and TAN (Figure 7G, H, J). Notably, lanifibranor prevented the TGF-β_1_-induced increase in NAD content (Figure 7K), resulting in a significantly lower ATP/NAD ratio (Figure 7L). Collectively, these data indicate that lanifibranor partially improves contractile function under TGF-β_1_ stimulation by normalizing contraction kinetics and beating rate, while largely preserving overall bioenergetic homeostasis despite measurable shifts in nucleotide balance.

### 2.5 Lanifibranor reversed the effects of TGF-β_1_ on the contractility of human cardiomyocytes cultured *in vitro*

The modulatory effects of lanifibranor on cardiac microtissue contractility prompted us to examine its impact on the contractile properties of human iPSC-derived cardiomyocytes (hiPSC-CMs). Using combined atomic force microscopy (AFM) and fluorescence imaging, we simultaneously assessed contraction force and calcium flux in hiPSC-CMs treated with lanifibranor (50 μM), in the absence or presence of TGF-β_1_ (10 ng/ml). TGF-β_1_ significantly increased contraction force and calcium flux, effects that were reversed by lanifibranor (Figure 8A-D). TGF-β_1_ also reduced beating rate, which was fully restored to near-control levels following lanifibranor treatment (Figure 8D). Mitochondrial respiration was assessed by measuring oxygen consumption rates (OCR) using the Seahorse XF Mito Stress Test (Figure 8E). TGF-β_1_ significantly increased basal and ATP-linked respiration without affecting maximal respiration or spare respiratory capacity. Lanifibranor normalized basal and ATP-linked respiration to near-control levels but did not reverse TGF-β_1_-induced changes in maximal respiration or spare respiratory capacity. Collectively, these findings indicate that lanifibranor improves cardiomyocyte contractile function under TGF-β_1_ stimulation not by enhancing mitochondrial respiratory capacity, but by improving the coupling between cellular energetics and contractile properties.

**Figure 8.**
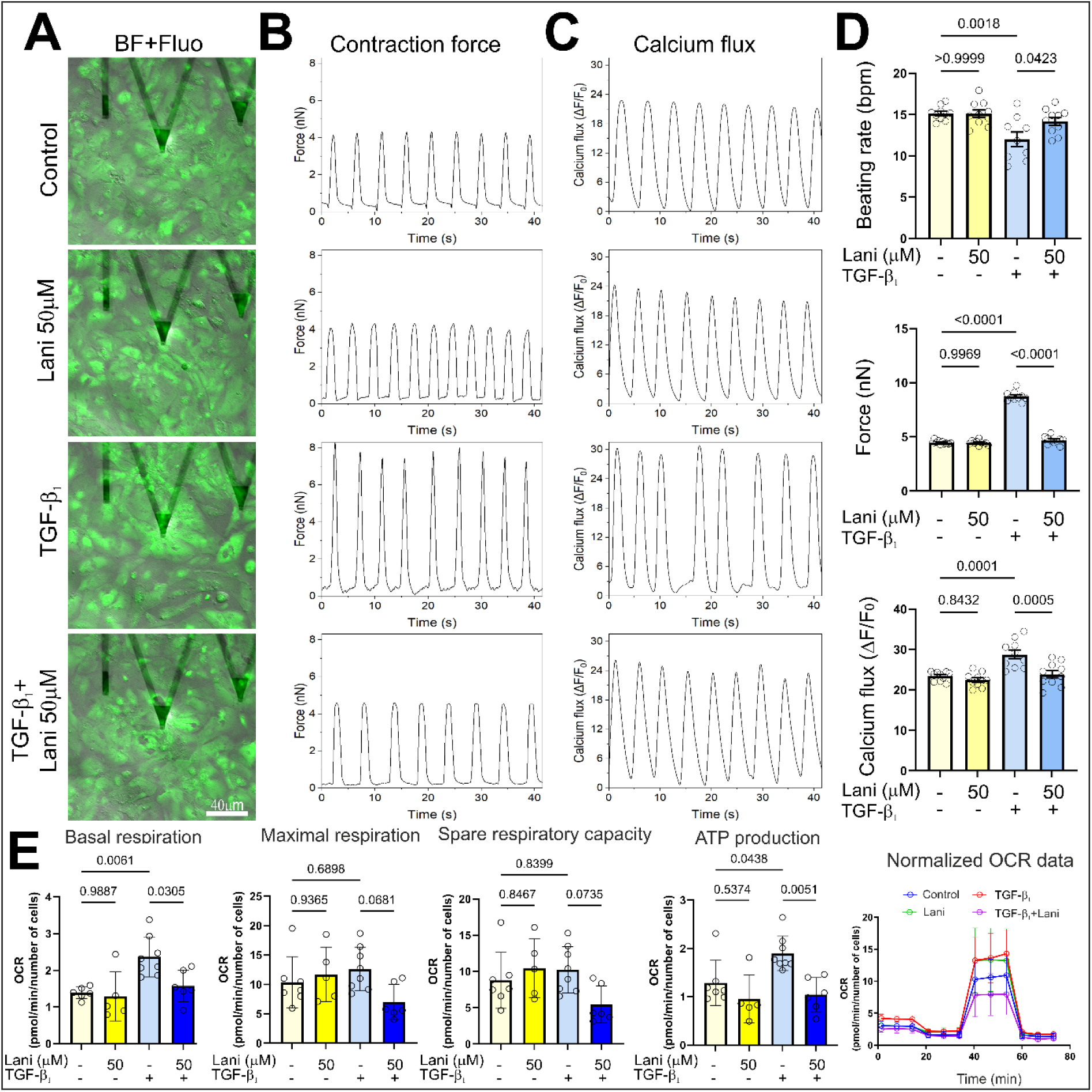
Effects of lanifibranor on contractility, calcium flux, and bioenergetics in TGF-β_1_-stimulated human iPSC-derived cardiomyocytes. Functional analysis of contractile properties and calcium flux was performed in human iPSC-derived cardiomyocytes (hiPSC-CMs) treated with lanifibranor (Lani; 50 μM) in the absence or presence of TGF-β_1_ (10 ng/ml), using combined atomic force microscopy (AFM) and fluorescence microscopy. **(A)** Representative merged bright-field (BF) and fluorescence (Fluo) images of hiPSC-CMs. **(B)** Representative contraction force traces and **(C)** calcium flux traces. **(D)** Quantitative analysis of functional parameters, including beating rate (bpm), contraction force, and calcium flux (n = 10). **(E)** Mitochondrial respiration assessed by oxygen consumption rate (OCR) measurements; OCR values were normalized to cell number and analyzed for basal respiration, maximal respiration, spare respiratory capacity, and ATP-linked respiration. Data are presented as mean ± SD. Statistical significance was determined using the Kruskal–Wallis test followed by Dunn’s post hoc test; p values are indicated in the graphs.

## 3. Discussion

Cardiac fibrosis represents a common pathological endpoint of diverse cardiovascular diseases, including hypertension, ischemic heart disease and cardiomyopathies, and is a major determinant of adverse clinical outcomes such as heart failure progression, arrhythmias, and increased mortality [32–34]. Despite extensive research, effective therapeutic modulation of cardiac fibrosis remains an unresolved clinical challenge [35]. Current pharmacological approaches primarily aim to slow down progression rather than reverse established fibrotic remodeling, largely due to the complex and highly interconnected signaling networks that govern extracellular matrix deposition and fibroblast activation [35, 36]. Among these, transforming growth factor-β (TGF-β) signaling plays a central role in driving myofibroblast differentiation and excessive matrix accumulation [26, 27, 32]. Although several experimental strategies targeting profibrotic pathways have shown promise in preclinical studies, their translation into routine clinical use has been limited by insufficient efficacy and safety concerns [35, 37]. Consequently, there is a growing need for improved experimental models that more accurately reflect the pathophysiological conditions of the human heart, enabling mechanistic studies and the identification of novel, more selective anti-fibrotic interventions [26, 27, 38].

In this study, we employed complementary *in vitro* models of increasing complexity, including two-dimensional (2D) cultures, three-dimensional (3D) constructs, and engineered cardiac microtissues, to investigate profibrotic cardiac remodeling. These human-relevant platforms enabled the evaluation of antifibrotic lanifibranor activity under controlled pro-fibrotic conditions and allowed assessment of its effects on fibroblast activation, extracellular matrix deposition, and tissue-level functional properties. Lanifibranor as a pan-peroxisome proliferator-activated receptor (PPAR) agonist has demonstrated clinical efficacy in randomized trials in liver fibrosis associated with nonalcoholic steatohepatitis (NASH/MASH) [39] and has shown robust anti-fibrotic effects across multiple preclinical models of fibrotic diseases, including liver, skin, and lung fibrosis [21–23, 40]. By integrating advanced 3D models with conventional 2D approaches, our work offers insights into the mechanisms by which lanifibranor may modulate profibrotic processes in the context of cardiac fibrosis.

The results presented in this study clearly demonstrate a strong TGF-β_1_-driven activation of cardiac fibroblasts across all tested experimental platforms, in line with previous reports. While fibroblasts are essential for maintaining structural integrity and synchronized contractile activity of engineered cardiac microtissues, their pathological activation leads to substantial changes in tissue architecture, ECM composition, and contractile properties [26, 41]. Lanifibranor consistently attenuated the fibroblast-to-myofibroblast phenotypic transition in 2D cultures, 3D spheroids, and multicellular cardiac microtissues. Transcriptomic and structural analyses indicated that lanifibranor modulates ECM-cell interaction pathways, particularly those related to integrin-dependent adhesion and signaling. Integrins link ECM composition and stiffness to cytoskeletal tension and sustained TGF-β signaling, thereby reinforcing the persistence of the myofibroblast phenotype [42]. In this context, the reduction of supermature focal adhesions - structures required for efficient transmission of α-SMA-dependent forces-suggests that lanifibranor may disrupt mechanotransduction feedback loops that stabilize the myofibroblast state even under persistent profibrotic stimulation [43, 44]. Importantly, previous studies have shown that PPAR agonists modulate focal adhesion–associated signaling, including integrin-FAK pathways, thereby influencing mechanotransduction in TGF-β_1_-driven fibroblast activation models [15, 45]. To our knowledge, the effects of a pan-PPAR agonist on mechanotransduction through modulation of cell-ECM interactions and focal adhesion remodeling have not previously been demonstrated in multicellular cardiac tissue models.

Pro-fibrotic TGF-β signaling disrupts contractile function in cardiac microtissues, leading to increased beating rate and impaired contraction-relaxation kinetics, as reported previously and confirmed in our study [26]. Importantly, lanifibranor counteracted these TGF-β_1_-induced alterations, preserving contractile parameters close to the non-fibrotic control state. To our knowledge, the effects of lanifibranor on cardiac contractile function have not previously been investigated *in vitro* or *in vivo*, in either basic or clinical studies. Our findings therefore provide the first evidence that lanifibranor-mediated inhibition of profibrotic remodeling is accompanied by improved contractile function in cardiac microtissues. Although lanifibranor has not been evaluated for direct cardiac functional outcomes, pharmacological activation of PPAR signaling has been shown to modulate myocardial contractility in preclinical models, including improved contractility and relaxation following treatment with the PPARα agonist AVE8134 in post-myocardial infarction rats and enhanced contractility induced by the PPARβ/δ agonist GW0742 in isolated hearts and diabetic rat models. Pharmacological activation of PPAR signaling has been shown to improve cardiac function through distinct, isoform-specific mechanisms, with PPARα agonist (AVE8134) primarily enhancing relaxation by limiting adverse remodeling and fibrosis, and PPARδ agonist (GW0742) directly increasing cardiomyocyte contractility via Ca²⁺-dependent modulation of the contractile apparatus [46, 47]. Together, these findings indicate that targeting cardiac fibrosis through PPAR-mediated pathways may have beneficial consequences also for tissue-level cardiac function.

Accumulating evidence indicates that fibroblast activation by TGF-β_1_ and their differentiation into myofibroblasts are accompanied by metabolic reprogramming, the direction of which depends on cell type, duration of stimulation, and functional context. In some models, TGF-β_1_ has been reported to increase mitochondrial respiration, which has been attributed to the higher energetic and biosynthetic demands of activated fibroblasts [48, 49]. In contrast, other studies describe a pattern in which TGF-β_1_ does not substantially affect basal respiration but reduces maximal respiration and spare respiratory capacity, indicating remodeling of mitochondrial reserve rather than a uniform decrease in metabolic activity [50, 51]. Our observations in hCFs are consistent with this latter scenario: TGF-β_1_ did not significantly alter basal respiration but reduced maximal respiration and spare respiratory capacity. This suggests that, in our system, FMT is accompanied by a shift in metabolic priorities rather than by simple suppression of cellular metabolism. Specifically, reduced respiratory reserve may reflect a transition away from a lipid-oxidative, mitochondria-supported metabolic program toward a profibrotic biosynthetic state associated with extracellular matrix production.

One possible explanation for divergent findings across models is the temporal and contextual nature of the TGF-β_1_ response, whereby early metabolic activation may evolve into a more specialized profibrotic state during sustained fibroblast activation and cell–ECM remodeling. In this context, the partial reversal of TGF-β_1_-associated reductions in maximal respiration and spare respiratory capacity by lanifibranor, together with increased ATP-linked respiration in hCFs, can be interpreted as restoration of bioenergetic adaptability accompanying attenuation of FMT. The nucleotide pool data obtained in cardiac microtissues do not necessarily contradict this interpretation, because end-point biochemical measurements in fixed microtissues reflect cellular energetic content rather than real-time metabolic flux.

Consistent with this interpretation, transcriptomic profiling of cardiac microtissues showed that TGF-β_1_ induced a predominantly profibrotic transcriptional program, with enrichment of extracellular matrix organization pathways and increased expression of matrix remodeling-and TGF-β-associated genes, including *TIMP3, COL11A1, COL8A2, PMEPA1, MXRA5, FAP, HAPLN1, MMP2, PTK7 and LTBP2*. In contrast, lanifibranor promoted metabolic and mitochondrial transcriptional signatures. In microtissues treated with lanifibranor alone, this was reflected by enrichment of pathways related to fatty acid metabolism, mitochondrial β-oxidation, lipid metabolism, cholesterol biosynthesis, steroid metabolism and respiratory electron transport, together with increased expression of metabolic genes such as *PDK4, ACSL1, HADHB, ANGPTL4, HADHA, HMGCS1* and *IDI1*. Under TGF-β_1_ stimulation, lanifibranor partially redirected the transcriptional response toward metabolic pathways, including fatty acid metabolism, cholesterol biosynthesis, respiratory electron transport and lipid metabolism by *PPARA, with increased expression of genes such as PDK4, ANGPTL4, UCP2, PLIN2, ABCA1, CDKN1A* and *PLPP1*. Thus, rather than indicating nonspecific metabolic recovery, lanifibranor appears to partially restore a PPAR-linked lipid-mitochondrial metabolic program while attenuating ECM-dominant profibrotic remodeling. This interpretation is further supported by structural data showing reduced α-SMA, fibronectin, periostin, collagen accumulation and procollagen 1α1 secretion in TGF-β_1_-treated microtissues. Together, available literature and our findings support the view that FMT and cardiac profibrotic remodeling are coupled to metabolic reprogramming, with lanifibranor acting as a pharmacological modulator that promotes PPAR-driven bioenergetic adaptation and weakens the myofibroblast/ECM program.

Available evidence indicates that TGF-β_1_ can directly influence cardiomyocyte function, although the direction and magnitude of these effects depend strongly on the experimental context. In isolated cardiomyocytes, TGF-β_1_ has been reported to impair contractile properties through disruption of excitation–contraction coupling, whereas contractile dysfunction is more consistently linked to fibroblast activation and extracellular matrix remodeling in multicellular cardiac tissue models [52, 53]. In our hiPSC-CM model, TGF-β_1_ increased contraction force and calcium transient amplitude while reducing beating rate, indicating a shift toward a stronger but slower contractile phenotype. These functional changes were accompanied by increased basal and ATP-linked respiration without alterations in maximal respiration or spare respiratory capacity, a profile consistent with elevated energetic demand per contraction rather than with impairment of mitochondrial function. Lanifibranor reversed the TGF-β_1_-induced alterations in contractile force, calcium handling, and beating rate, while normalizing basal respiration and ATP production toward control levels without increasing mitochondrial respiratory reserve. While direct cardiomyocyte-specific data on lanifibranor from independent studies remain limited, other PPAR agonists have shown cardioprotective effects in preclinical models, including improvements in contractility and calcium signaling [46, 47]. Our findings support the conclusion that lanifibranor does not exert deleterious effects on cardiomyocytes but instead restores physiological contractile properties, contributing to a favorable safety and efficacy profile in cardiac disease associated with fibrosis.

In conclusion, rather than globally suppressing TGF-β signaling, lanifibranor appears to modulate downstream programs related to cell adhesion, cytoskeletal tension, and metabolism, thereby promoting maintenance of cardiac fibroblasts in a non-activated state under persistent profibrotic stimulation. To our knowledge, this is the first study demonstrating that lanifibranor attenuates fibrotic processes in human cardiac models under TGF-β_1_-driven conditions. The use of complementary 2D and 3D human-based platforms, combined with integrated molecular, structural, and functional readouts, enabled a comprehensive characterization of cardiac fibroblast phenotype regulation, cardiomyocyte responses, and multicellular cardiac microtissue behavior.

Several limitations should be considered when interpreting these findings. Cardiac fibrosis was modeled primarily through TGF-β_1_ stimulation, which captures a key profibrotic pathway but does not fully recapitulate the complexity of in vivo disease. In addition, lanifibranor was administered concomitantly with TGF-β_1_; therefore, the observed effects predominantly reflect attenuation of ongoing fibrotic remodeling rather than reversal of established fibrosis. The potential involvement of mechanotransduction in these processes warrants further investigation, particularly regarding the interplay between extracellular matrix mechanical properties and YAP/TAZ and integrin/FAK signaling pathways.

Despite these limitations, our integrated experimental approach demonstrates coordinated changes at the transcriptomic, extracellular matrix, and tissue-functional levels during lanifibranor-mediated attenuation of profibrotic responses in cardiac tissue. Together with prior clinical evidence demonstrating antifibrotic efficacy and a favorable safety profile of lanifibranor in fibrometabolic diseases [23, 39], these findings support continued investigation of lanifibranor and broader PPAR-targeted strategies for the treatment and prevention of fibrosis-driven cardiomyopathies

## 4. Materials and methods

### 4.1 Cell cultures and microtissue formation

In this study, fetal and adult human cardiac fibroblasts (hCFs; Cell Applications/Merck, cat. no. 306-05F and 306-05A, respectively) were used. Cells were cultured under standard conditions (37 °C, 5% CO₂) in Dulbecco’s Modified Eagle Medium with high glucose (DMEM-HG; Sigma-Aldrich/Merck), supplemented with 10% fetal bovine serum (FBS; Gibco) and a penicillin/streptomycin cocktail (Gibco). For 2D experiments, hCFs between passages 6 and 15 were seeded at a density of 5 000 cells/cm². After 24 hours, the culture medium was replaced by Maintenance Medium (MM; DMEM-HG supplemented with 2% FBS, 50 μM phenylephrine hydrochloride (Sigma-Aldrich/Merck), 0.3 μM L-ascorbic acid (Sigma-Aldrich/Merck), and 50 U/ml penicillin/streptomycin (Gibco) with or without lanifibranor (Cayman Chemical; 0 - 125 μM). A profibrotic response was induced by the addition of recombinant human TGF-β_1_ (PeproTech; 10 ng/ml). Cells were cultured under these conditions for 4 days. To the formation of 3D cultures of hCFs, 5 000 cells were suspended in 80 μl of MM and seeded into each well of 96-well GravityTRAP plates (InSphero, Schlieren, Switzerland). 3D hCFs cultures were incubated under standard conditions (37 °C, 5% CO₂) in a tilted position for 2 days to facilitate self-assembly and then the medium was replaced by fresh MM without or with tested compounds. Spheroids were maintained for 10 days, with medium changes performed every 2 days.

Human cardiomyocytes (hiPSC-CMs) derived from induced pluripotent stem cells (iPSCs) were obtained either from FujiFilm Cellular Dynamics (iCell® Cardiomyocytes2, cat. #01434) or generated from the episomal hiPSC line (Gibco, A18945) as previously described [54]. For 2D experiments, hiPSC-CMs were seeded on Geltrex-coated plates (LDEV-free, reduced growth factor basement membrane matrix; Gibco) in RPMI-1640 medium supplemented with 2% B27 with insulin (Gibco). Metabolic selection using 4 mM sodium lactate (Sigma-Aldrich/Merck) in glucose-free DMEM (Gibco) was performed to obtain purified cardiomyocyte cultures. Cells were stimulated with lanifibranor (50 μM), with or without TGF-β_1_ (10 ng/ml), and cultured under standard conditions (37 °C, 5% CO₂) for 24 hours.

To microtissues (MTs) assembly co-cultures of hiPSC-CMs and hCFs were used. Cells were mixed at a 4:1 ratio (hiPSC-CMs:hCFs, 5000 cells/spheroid), suspended in 80 μl of MM and seeded into each well of 96-well GravityTRAP plates (InSphero, Schlieren, Switzerland). Cultures were incubated under standard conditions (37 °C, 5% CO₂) in a tilted position for 2 days to facilitate self-assembly. Then, the medium was replaced with fresh MM containing lanifibranor (0 - 125 μM) in the absence or presence of TGF-β_1_ (20 ng/ml). Higher TGF-β_1_ concentrations in 3D models compared with 2D cultures were used due to limited diffusion, which is in line with previous reports [26]. MTs were cultured for 10 days, with medium changes performed every 2 days.

### 4.2 Evaluation of contractile function of microtissues and hiPSC-CMs

Contractility of human cardiomyocytes (hiPSC-CMs) was assessed using a combined atomic force microscopy (AFM) and fluorescence microscopy system (BioScope Catalyst AFM mounted on a Zeiss inverted optical microscope), equipped with an ORCA-Flash4.0 LT3 digital CMOS camera. Briefly, Fluo-4 (Invitrogen) was added to the culture dish and incubated with the cells for 30 minutes. After positioning the cells in the center of the AFM’s optical path (at 37 °C), 10 cells were selected for each experimental condition. Measurements were performed using an MLCT-Bio probe with a nominal spring constant of 0.01 N/m. The AFM was activated to establish contact with each individual cell. Simultaneous acquisition of AFM and fluorescence images was conducted using a custom-made script. The same camera and image acquisition settings were applied throughout the fluorescence image collection. Raw AFM data were converted into force values using a calibration file and custom-developed software. Two-step calibration (pre-and post-measurement) was performed, and averaged values were used to calculate beating rate and contraction force. Calcium flux was quantified using a dedicated ImageJ plugin.

Microtissue contractility was assessed using the AxioObserver Z1 microscope (Zeiss, Hombrechtikon, Switzerland) and ZEN software. Captured images were converted into videos using Fiji (ImageJ) software and a custom-made macro. Contractile properties of the microtissues were then analyzed using Fiji and the MUSCLEMOTION macro [31].

### 4.3 Cell viability and metabolic activity assays

Cell metabolic activity and viability was determined following exposure of cells/spheroids to increasing concentrations of lanifibranor (0 - 125 μM) in the absence or presence of TGF-β_1_ (10 ng/ml for 2D cultures and 20 ng/ml for 3D cultures). Two metabolic activity-based assays based on the PrestoBlue™ HS Cell Viability Reagent (Invitrogen) and MTT were performed in this study according to the manufacturer’s instructions. The PrestoBlue reagent were diluted at a 10:1 ratio in fresh MM, added to each well containing cells or spheroids and incubated for 4 hours at 37 °C and 5% CO₂. In case of MTT assay, MTT reagent (5 mg/ml; Sigma-Aldrich/Merck) was added to conditioned medium (final concentration 0.5 mg/ml) and incubated with cells for 4 hours under standard conditions (37 °C, 5% CO₂). The medium was then replaced and 100 μl of isopropyl alcohol was added to each well to solubilize the formazan product. The fluorescence of resorufin was measured at 590 nm and absorbance of solubilized formazan was measured at 570 nm using a Synergy (BioTek) with Gen5 software or Multiskan FC (Thermo Fisher Scientific) microplate reader.

Cell viability was assessed using the fluorescein diacetate (FDA)/ethidium bromide (EtBr) assay (Sigma-Aldrich/Merck). Following treatment, cells were trypsinized and resuspended in phosphate-buffered saline (PBS) containing FDA/EtBr solution. Viable (FDA⁺/EtBr⁻) and non-viable (EtBr⁺) cells were counted using a Leica DMI6000B fluorescence microscope equipped with LAS X software (Leica Microsystems GmbH, Wetzlar, Germany). Results were expressed as the percentage of viable cells under each tested condition (25 - 125 μM) relative to the untreated control (0 μM).

### 4.4 Caspase 3/7 activity assay

Microtissues (MTs) were transferred into fresh medium in white-walled 96-well plates, and an equal volume of Caspase-Glo® 3/7 reagent (Promega, Dübendorf, Switzerland) was added to each well. After gentle mixing, the plates were incubated at room temperature for 3 hours, protected from light. Luminescence was then measured using a Synergy microplate reader (BioTek, Winooski, Vermont, USA) and analyzed with Gen5 software.

### 4.5 á-SMA and procollagen 1α1 quantification by ELISA

Human cardiac fibroblasts (hCFs) cultured under the 2D conditions described above for 4 days were fixed and permeabilized with ice-cold methanol, then blocked with 1% BSA in PBS containing 0.1% Tween-20 for 60 minutes. Cells were incubated overnight at 4 °C with a primary antibody against α-SMA (mouse monoclonal anti-α-SMA; Sigma-Aldrich/Merck) diluted in 1% BSA/PBS. After three washes, cells were incubated with goat anti-mouse secondary antibodies conjugated to horseradish peroxidase (HRP) for 1 hour at room temperature in 1% BSA/PBS. Tetramethylbenzidine (TMB; Sigma-Aldrich/Merck) was used to initiate the colorimetric reaction, which was stopped with 1N HCl. Absorbance was measured at 450 nm using a Synergy microplate reader (BioTek, Winooski, VT, USA) or a Multiskan FC reader (Thermo Fisher Scientific).

Supernatants from 2D hCF cultures (day 4) and from individual 3D microtissue cultures (day 10) were collected and stored at-80 °C. The concentration of procollagen 1α1 was measured using the Human Procollagen I Alpha 1 DuoSet ELISA kit (R&D Systems, Abingdon, UK) according to the manufacturer’s instructions. Absorbance was read at 450 nm using the Synergy microplate reader (BioTek, Winooski, VT, USA) and analyzed with Gen5 software.

### 4.6 Immunofluorescence and histochemical staining and image analysis

Cells were seeded on glass coverslips in a 12-well plate and cultured under the conditions described above for 4 days. Subsequently, cells were fixed and immunostained according to a previously published protocol [15]. The following primary antibodies were used: mouse monoclonal IgG anti-α-SMA, mouse monoclonal IgG anti-vinculin, and rabbit polyclonal IgG anti-collagen I (all from Sigma-Aldrich/Merck). Appropriate secondary antibodies conjugated to AlexaFluor 488 or AlexaFluor 546 (goat anti-mouse or goat anti-rabbit; Life Technologies, Thermo Fisher Scientific, Waltham, MA, USA) were applied. Hoechst 33258 (1 µg/ml; Sigma-Aldrich/Merck) was used for staining of cell nucleus. F-actin bundles were visualized using AlexaFluor 546–conjugated phalloidin (Life Technologies, Thermo Fisher Scientific). Images were acquired using a Leica DMI6000B fluorescence microscope equipped with a TIRF (Total Internal Reflection Fluorescence) module and LasX software (v3.7.4; Leica Microsystems GmbH, Wetzlar, Germany). The area and length of focal adhesions were quantified using a custom-made macro developed for Fiji (ImageJ; NIH, Bethesda, MD, USA).

For immunohistochemical analysis, 25–30 cardiac microtissues were pooled, fixed overnight in 4% paraformaldehyde in PBS and embedded in 1% agarose. Samples were then dehydrated through graded ethanol, cleared in xylene, and embedded in paraffin at 56°C. Paraffin sections of 2 µm thickness were mounted on Superfrost Plus slides and dried overnight at 58°C. Sections were deparaffinized in xylene, rehydrated through descending ethanol concentrations, washed in distilled water, and subjected to antigen retrieval in Tris-EDTA buffer, pH 9, or citrate buffer, pH 6. After blocking with 10% goat serum (Vector Laboratories), sections were incubated with primary antibodies against α-SMA (Abcam ab132575, clone E184; 1:2000), periostin (Abcam ab14041; 1:2000), fibronectin (Abcam ab2413; 1:2000), and troponin T (Invitrogen MA4-12960; 1:20000), followed by detection using the Bond Polymer HRP Refine Detection Kit (Leica). Nuclei were counterstained with haematoxylin. Collagen content was assessed by Masson’s Trichrome staining. Images were acquired using a Leica DMi8 microscope equipped with CoolLED pE-4000 illumination and AFC autofocus stabilization. Immunoreactive signal and nuclear staining were quantified in Fiji, and signal intensity was normalized to the number of counterstained nuclei and reported as arbitrary units (AU).

### 4.7 Intracellular nucleotide quantification

Microtissues (MTs) were suspended in 100 μl of 80% methanol and stored at −80 °C. Sample preparation and analysis of intracellular nucleotide content were performed using ultra-high-performance liquid chromatography with diode array detection (UHPLC-DAD), using the Nexera LC-40 system and an SPD-M30A diode array detector equipped with a high-sensitivity 85-mm optical path cell (Shimadzu, Japan), according to a previously described protocol [55]. The concentrations of AMP, ADP, ATP, NAD were expressed in nanomoles per gram of dry tissue (nmol/g). The ATP/NAD ratio were calculated, Adenylate energy charge (AEC) was calculated according to the Atkinson definition as (ATP + 0.5 × ADP)/(ATP + ADP + AMP), and the total adenine nucleotide pool (TAN) was defined as the sum of ATP, ADP, and AMP, using intracellular nucleotide concentrations measured by UHPLC-DAD.

### 4.8 Mitochondrial respiration analysis (Seahorse XF assay)

hCFs or hiPSC-CMs were seeded into Seahorse XFe96 plates at 5000 or 15000 per well, respectively, in dedicated complete medium. After 24 h of incubation, the medium was replaced by fresh medium supplemented with lanifibranor without or with TGF-β_1_ and cultured for 4 days. The Seahorse analyzer (Seahorse Bioscience, N. Billerica, MA, USA) were used to the examine of cellular respiration parameters with a Agilent Seahorse XF Cell Mito Stress Test Kit. Before measurement, the medium was replaced by bicarbonate-free Agilent Seahorse XF Base Medium Minimal Dulbecco’s modified Eagle’s medium (DMEM) or DMEM (D5030, Sigma-Aldrich) supplemented with 10 mM glucose, 2 mM L-glutamine, 1 mM sodium pyruvate (pH 7.4). Cultured cells were washed twice with a suitable assay medium and incubated for 45 min at 37 °C without CO_2_. Concentrations of standard mitochondrial modulators were optimized in preliminary experiments for both types of cells separately. The mitochondrial stress test was performed by sequential injection of oligomycin (ATP synthase inhibitor, 1 μg/ml), carbonyl cyanide-p-trifluoromethoxyphenylhydrazone (FCCP; uncoupler, 9 μM or 2 μM for hCFs or hiPSC-CMs, respectively), and a combination of rotenone and antimycin A (complex I and III inhibitors, 0.5 μM for each compound). Basal respiration, ATP-linked respiration, maximal respiration, and spare respiratory capacity were calculated from OCR traces using Seahorse Wave software. OCR values were normalized to cell number or total protein content, as indicated. Each experimental condition was measured in multiple technical and three independent biological replicates.

### 4.9 RNA sequencing and data analysis

RNA extraction, library preparation, sequencing, and primary bioinformatic processing were performed by Lexogen. RNA-seq libraries were prepared using Lexogen’s LUTHOR High-Definition 3′ mRNA-seq protocol for ultra-low input RNA samples, incorporating unique molecular identifiers (UMIs) for gene-level deduplication. Libraries were sequenced on an Illumina platform in paired-end 2 × 100 bp mode, yielding approximately 5 million reads per sample on average (range: 3-8 million). Reads were trimmed with cutadapt, quality-controlled using FastQC and MultiQC, and aligned to the human reference genome GRCh38 with STAR. Gene-level quantification, UMI-based deduplication, and differential expression analysis were performed using DESeq2. Differentially expressed genes were defined as genes with |log₂ fold change| ≥ 1 and adjusted p-value < 0.05. Gene set enrichment analysis (GSEA) was performed using preranked gene lists based on the DESeq2 Wald statistic and the fgsea package, with Hallmark, Reactome, and GO Biological Process gene sets obtained from msigdbr. Gene sets containing 15–500 genes were tested, and pathways with FDR < 0.05 were considered significantly enriched. Reactome GSEA was used for the main pathway-level interpretation. For supplementary analyses, Reactome over-representation analysis (ORA) was performed separately for upregulated and downregulated DEGs using clusterProfiler::enricher, with all genes from the corresponding DESeq2 result table used as the background universe. Reactome ORA gene sets containing 10 - 500 genes were tested, and FDR < 0.05 was used as the significance threshold. Data visualization and downstream analyses were performed using custom R scripts. RNA sequencing data generated in this study will be deposited in the Gene Expression Omnibus and accession numbers will be provided before publication.

### 4.10 Statistical Analysis

All quantitative data are presented as mean ± standard deviation (SD). The normality of data distribution was assessed using the Shapiro–Wilk test. Depending on the distribution, statistical significance was determined using either the nonparametric Kruskal–Wallis test followed by Dunn’s multiple comparisons post hoc test, or one-way analysis of variance (ANOVA) with Tukey’s multiple comparisons post hoc test. Individual p values are indicated in each graph. All statistical analyses were performed using GraphPad Prism version 10.4.2.

## Acknowledgements

The authors thank the Center for Microscopy and Image Analysis of the University of Zurich for technical assistance in data acquisition. Graphical abstract was created in BioRender. Paw, M. (2026) https://BioRender.com/q5cqzm5.

## Author Contributions

Conceptualization, M.P., G.K.;

Methodology, M.P., L.M., A.L., P.K., B.K.-Z., A.B., M.S., S.B.-W., D.W., M.C.;

Software, M.P., L.M.; Validation, M.P.; Formal Analysis, M.P.; Investigation, M.P.;

Resources, M.P., P.B. and G.K.; Data Curation, M.P.;

Writing—Original Draft Preparation, M.P.;

Writing—Review & Editing, L.M., A.L., M.C., B.K.-Z., A.B., M.S., S.B.-W., D.W., P.K., S.C., P.B., Z.M., O.D., J.C. G.K. and M.P.;

Visualization, M.P.; Supervision, J.C. and G.K.;

Project Administration, M.P. and G.K.;

Funding Acquisition, M.P., P.B., Z.M., O.D. and G.K.

All authors have read and agreed to the published version of the manuscript.

## Ethics approval

Not applicable.

## Consent to participate

Not applicable.

## Data Availability Statement

RNA sequencing data generated in this study will be deposited in a public gene expression repository (Gene Expression Omnibus, GEO) and will be made available upon publication. Accession numbers will be provided in the final version of the manuscript. All other data supporting the findings of this study will be available from the corresponding author upon reasonable request.

## Declaration of generative AI and AI-assisted technologies in the manuscript preparation process

During the preparation of this work, the authors used ChatGPT by OpenAI to assist with language editing and text refinement. After using this tool, the authors reviewed and edited the content as needed and take full responsibility for the content of the published article. No generative AI or AI-assisted tools were used to create or alter figures, images, artwork, experimental data, or scientific results.

## Abbreviations

α-SMA: α-smooth muscle actin
AEC: adenylate energy charge
AFM: atomic force microscopy
AMP: adenosine monophosphate
ANOVA: analysis of variance
AP: alkaline phosphatase
ATP: adenosine triphosphate
AU: arbitrary units
BSA: bovine serum albumin
CVDs: cardiovascular diseases
DAB: 3,3′-diaminobenzidine
DEGs: differentially expressed genes
DMEM-HG: Dulbecco’s Modified Eagle Medium (high glucose)
ECM: extracellular matrix
ELISA: enzyme-linked immunosorbent assay
EtBr: ethidium bromide
FA(s): focal adhesion(s)
FBS: fetal bovine serum
FDA: fluorescein diacetate (viability dye)
FMT: fibroblast-to-myofibroblast transition
GEO: Gene Expression Omnibus
hiPSC(s): human induced pluripotent stem cell(s)
hCF(s): human cardiac fibroblast(s)
hCM(s): human cardiomyocyte(s)
HRP: horseradish peroxidase
IL-1β: interleukin 1 beta
iPSC(s): induced pluripotent stem cell(s)
MM: maintenance medium
MT(s): microtissue(s)
MTT: 3-(4,5-dimethylthiazol-2-yl)-2,5-diphenyltetrazolium bromide
NAD: nicotinamide adenine dinucleotide
NASH: nonalcoholic steatohepatitis
OCR: oxygen consumption rate
PBS: phosphate-buffered saline
PCA: principal component analysis
PPAR(s): peroxisome proliferator-activated receptor(s)
SD: standard deviation
SEM: standard error of the mean
TAN: total adenine nucleotide pool
TGF-β /: TGF-β_1_ transforming growth factor beta / beta 1
UHPLC-DAD: ultra-high-performance liquid chromatography with diode array detection

## Supplementary data

**Figure S1.**
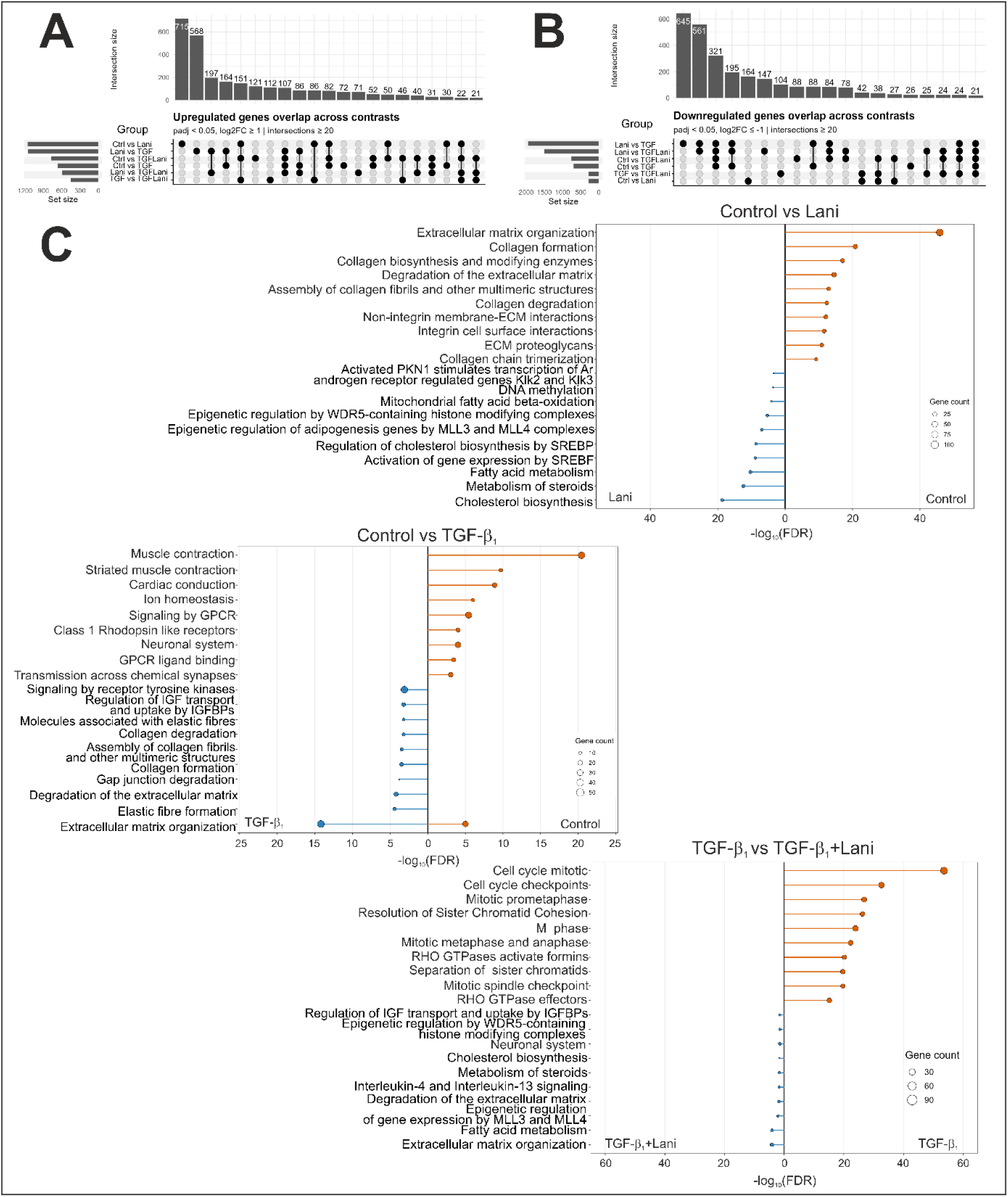
Reactome ORA of DEG sets across pairwise comparisons. **(A-B)** UpSet plots showing the overlap of upregulated **(A)** and downregulated **(B)** DEGs across contrasts. **(C)** Reactome over-representation analysis performed separately for DEG sets upregulated in each direction of the indicated comparisons. Paired lollipop plots show significantly enriched Reactome pathways plotted toward the condition in which the corresponding DEG set was upregulated; the x-axis represents mirrored-log₁₀(FDR), and point size denotes the number of DEGs associated with each pathway. DEGs were defined as |log₂FC| ≥ 1 and adjusted p-value < 0.05. ORA highlighted lipid/cholesterol metabolism, mitochondrial fatty acid β-oxidation, steroid metabolism, and epigenetic pathways among lanifibranor-upregulated DEGs; ECM/collagen remodeling, elastic fiber organization, gap junction degradation, and IGF/IGFBP-related pathways among TGF-β₁-upregulated DEGs; and ECM organization, fatty acid and cholesterol metabolism, IL-4/IL-13 signaling, IGF/IGFBP-related processes, and epigenetic regulation among TGF-β₁ + lanifibranor-upregulated DEGs. Opposite-side enrichments included ECM/collagen and integrin-related pathways in control-upregulated DEGs versus lanifibranor, contractile/cardiac conduction and ion-homeostasis pathways in control-upregulated DEGs versus TGF-β₁, and cell cycle, mitotic checkpoint, chromatid cohesion/segregation, and RHO GTPase-related pathways in TGF-β₁-upregulated DEGs versus TGF-β₁ + lanifibranor.

---

**Table S1.**
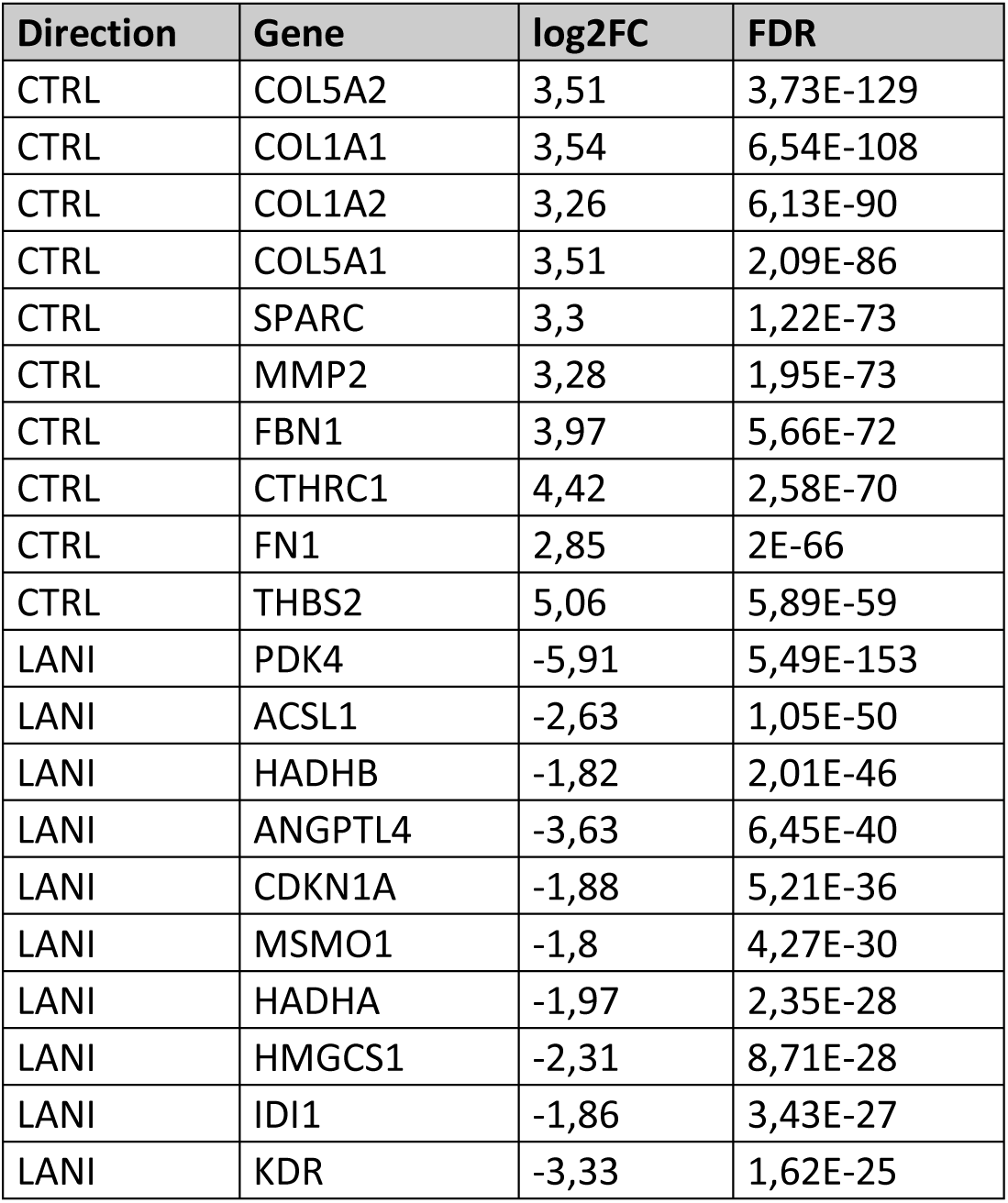
List of top differentially expressed genes in Control (CTRL) vs lanifibranor (LANI) comparison. Differentially expressed genes (DEGs) were defined as genes with FDR < 0.05 and |log2FC| ≥ 1. The table shows the top 10 genes upregulated in Control and the top 10 genes upregulated in lanifibranor, selected from the DEG list using a predefined ranking procedure. Positive log2FC values indicate higher expression in Control, whereas negative log2FC values indicate higher expression in lanifibranor.

| Direction | Gene | log<sub>2</sub>FC | FDR |
| --- | --- | --- | --- |
| CTRL | COL5A2 | 3,51 | 3,73E-129 |
| CTRL | COL1A1 | 3,54 | 6,54E-108 |
| CTRL | COL1A2 | 3,26 | 6,13E-90 |
| CTRL | COL5A1 | 3,51 | 2,09E-86 |
| CTRL | SPARC | 3,3 | 1,22E-73 |
| CTRL | MMP2 | 3,28 | 1,95E-73 |
| CTRL | FBN1 | 3,97 | 5,66E-72 |
| CTRL | CTHRC1 | 4,42 | 2,58E-70 |
| CTRL | FN1 | 2,85 | 2E-66 |
| CTRL | THBS2 | 5,06 | 5,89E-59 |
| LANI | PDK4 | -5,91 | 5,49E-153 |
| LANI | ACSL1 | -2,63 | 1,05E-50 |
| LANI | HADHB | -1,82 | 2,01E-46 |
| LANI | ANGPTL4 | -3,63 | 6,45E-40 |
| LANI | CDKN1A | -1,88 | 5,21E-36 |
| LANI | MSMO1 | -1,8 | 4,27E-30 |
| LANI | HADHA | -1,97 | 2,35E-28 |
| LANI | HMGCS1 | -2,31 | 8,71E-28 |
| LANI | IDI1 | -1,86 | 3,43E-27 |
| LANI | KDR | -3,33 | 1,62E-25 |

**Table S2.** List of top differentially expressed genes in Control (CTRL) vs TGF-β_1_ (TGF-β) comparison. Differentially expressed genes (DEGs) were defined as genes with FDR < 0.05 and |log2FC| ≥ 1. The table shows the top 10 genes upregulated in Control and the top 10 genes upregulated in lanifibranor, selected from the DEG list using a predefined ranking procedure. Positive log2FC values indicate higher expression in Control, whereas negative log2FC values indicate higher expression in TGF-β_1_.

| Direction | Gene | log2FC | FDR |
| --- | --- | --- | --- |
| CTRL | CHI3L1 | 2,7 | 2,19E-44 |
| CTRL | C1orf105 | 2,36 | 1,24E-30 |
| CTRL | BNIP3 | 2,34 | 1,37E-26 |
| CTRL | GPM6B | 3,9 | 1,8E-23 |
| CTRL | ANK3 | 1,8 | 1,1E-21 |
| CTRL | NAMPT | 1,5 | 2,53E-21 |
| CTRL | STC2 | 2,29 | 5,77E-21 |
| CTRL | RPS6KA2 | 1,5 | 1,2E-20 |
| CTRL | BMP5 | 1,57 | 1,08E-19 |
| CTRL | SAT1 | 1,58 | 1,26E-19 |
| TGF-β | TIMP3 | -3,31 | 8,29E-123 |
| TGF-β | COL11A1 | -3,71 | 1,06E-73 |
| TGF-β | COL8A2 | -3,82 | 1,42E-64 |
| TGF-β | PMEPA1 | -2,61 | 6,81E-59 |
| TGF-β | MXRA5 | -2,71 | 4,57E-55 |
| TGF-β | FAP | -2,62 | 4,33E-48 |
| TGF-β | HAPLN1 | -4,84 | 9,28E-48 |
| TGF-β | MMP2 | -2,15 | 1,34E-47 |
| TGF-β | PTK7 | -2,61 | 1,44E-47 |
| TGF-β | LTBP2 | -2,86 | 1,35E-46 |

**Table S3.** List of top differentially expressed genes in TGF-β_1_ (TGF-β) vs lanifibranor+TGF-β_1_ (LANI+TGF-β) comparison. Differentially expressed genes (DEGs) were defined as genes with FDR < 0.05 and |log2FC| ≥ 1. The table shows the top 10 genes upregulated in Control and the top 10 genes upregulated in lanifibranor, selected from the DEG list using a predefined ranking procedure. Positive log2FC values indicate higher expression in Control, whereas negative log2FC values indicate higher expression in TGF-β_1_.

| Direction | Gene | $\log_2FC$ | FDR |
| --- | --- | --- | --- |
| TGF-β | TOP2A | 5,2 | 3,69E-27 |
| TGF-β | TMEM158 | 4,47 | 9,14E-27 |
| TGF-β | STMN1 | 1,79 | 9,07E-26 |
| TGF-β | PTTG1 | 3,14 | 9,42E-23 |
| TGF-β | ADAM19 | 2,01 | 9,42E-23 |
| TGF-β | ANLN | 3,5 | 1,06E-22 |
| TGF-β | PCLAF | 4,46 | 4,76E-22 |
| TGF-β | UBE2C | 5,47 | 1,56E-20 |
| TGF-β | AQP1 | 2,54 | 3,17E-20 |
| TGF-β | RRM2 | 4,34 | 5,13E-20 |
| LANI+TGF-β | PDK4 | -5,15 | 2,85E-102 |
| LANI+TGF-β | CHI3L1 | -3,75 | 4,56E-88 |
| LANI+TGF-β | ANGPTL4 | -6,59 | 2,61E-85 |
| LANI+TGF-β | UCP2 | -2,67 | 4,85E-35 |
| LANI+TGF-β | PLIN2 | -2,42 | 6,35E-35 |
| LANI+TGF-β | ABCA1 | -2,6 | 3,4E-33 |
| LANI+TGF-β | CDKN1A | -1,81 | 1,17E-32 |
| LANI+TGF-β | PLPP1 | -1,83 | 3,21E-25 |
| LANI+TGF-β | NEAT1 | -1,18 | 1,75E-21 |
| LANI+TGF-β | COMP | -2,38 | 2,64E-19 |

